# Scots pine – panmixia and the elusive signal of genetic adaptation

**DOI:** 10.1101/2023.06.09.543371

**Authors:** Jade Bruxaux, Wei Zhao, David Hall, Alexandru Lucian Curtu, Piotr Androsiuk, Andreas D. Drouzas, Oliver Gailing, Heino Konrad, Alexis R. Sullivan, Vladimir Semerikov, Xiao-Ru Wang

**Affiliations:** Department of Ecology and Environmental Science, Umeå Plant Science Center, Umeå University, Umeå, Sweden; Forestry Research Institute of Sweden (Skogforsk), Sävar, Sweden; Department of Silviculture, Transilvania University of Braşov, Braşov, Romania; Department of Plant Physiology, Genetics and Biotechnology, Faculty of Biology and Biotechnology, University of Warmia and Mazury in Olsztyn, Olsztyn, Poland; Laboratory of Systematic Botany and Phytogeography, School of Biology, Aristotle University of Thessaloniki, Thessaloniki, Greece; Department of Forest Genetics and Forest Tree Breeding, University of Göttingen, Göttingen, Germany; Austrian Research Centre for Forests (BFW), Department of Forest Biodiversity and Nature Conservation, Unit of Ecological Genetics, Vienna, Austria; Institute of Plant and Animal Ecology, Ural Division of Russian Academy of Sciences, Ekaterinburg, Russia

**Author notes:** These authors contributed equally to the work (Jade Bruxaux, Wei Zhao, David Hall). Correspondence to Xiao-Ru Wang.

**Keywords:** Conifer, demography, genetic diversity, gene flow, genetic-environmental association, population structure, *Pinus sylvestris*

## Abstract

- Scots pine is the foundation species of diverse forested ecosystems across Eurasia and displays remarkable ecological breadth, occurring in environments ranging from temperate rainforests to arid tundra margins. Such expansive distributions can be favored by various demographic and adaptive processes and the interactions between them.
- To understand the impact of neutral and selective forces on genetic structure in Scots pine, we conducted range-wide population genetic analyses on 2,321 trees from 202 populations using genotyping-by-sequencing, reconstructed the recent demography of the species, and examined signals of genetic adaptation.
- We found a high and uniform genetic diversity across the entire range (global *F*_ST_ 0.048), no increased genetic load in expending populations and minor impact of the last glacial maximum on historical population sizes. Genetic-environmental associations identified only a handful of SNPs significantly linked to environmental gradients.
- The results suggest that extensive gene flow is predominantly responsible for the observed genetic patterns in Scots pine. The apparent missing signal of genetic adaptation is likely attributed to the intricate genetic architecture controlling adaptation to multi-dimensional environments. The panmixia metapopulation of Scots pine offers a good study system for further exploration into how genetic adaptation and plasticity evolve under gene flow and changing environment.

## Introduction

Life history traits, demographic events, and selection shape the amount and distribution of genetic diversity across a species range (Ellegren & Galtier, 2016). Wind-pollinated outcrossing species are generally expected to have higher genetic diversity and weaker genetic structure compared to inbreeding species. In a heterogeneous fitness landscape, this high genetic diversity would provide a greater adaptive potential and promote divergence under environment-specific selection (Hamrick & Godt, 1996; Savolainen *et al*., 2013; Gamba & Muchhala, 2020). Consecutive range expansions after contraction may carry only a portion of the diversity from the source population and produce differentiation via genetic drift (Hewitt, 1996; Orsini *et al*., 2013; Wang & Bradburd, 2014; Zhao *et al*., 2020). These processes could potentially constrain a species ability to track, tolerate, and adapt to changing environments through their impact on standing adaptive genetic diversity or on gene flow between populations. Understanding the legacy effect of these processes is particularly important for boreal forest trees, which are subjected to strong ongoing climate change (Davis & Shaw, 2001; Koenigk *et al*., 2020).

Scots pine (*Pinus sylvestris* L.) is the foundation species of diverse forested ecosystems across Eurasia. It has the widest distribution of any pine species, with a range spanning from the Iberian and Balkan Peninsulas north to the Barents Sea, and from Britain and Ireland east to the Russian Pacific coast (Fig. **1**; Carlisle & Brown, 1968). It has an exceptionally wide niche, tolerating annual average temperatures ranging from -3°C to 16°C, and annual precipitation between 400 and 2,900 mm (Houston Durrant *et al*., 2016). It can be found from the sea level in its most northern populations and up to 2,600 m above the sea level in the Caucasus (Houston Durrant *et al*., 2016). It is a light demanding species able to grow on very nutrient poor sites (Houston Durrant *et al*., 2016). The variable habitat types that Scots pine occupies appear to have promoted distinct local adaptation as reflected by characteristic growth forms, ecotypes and clines in physiological traits, with up to 150 morphological varieties described (Carlisle & Brown, 1968; Rehfeldt *et al*., 2002; Tóth *et al*., 2017a). Phenotypic differences on traits such as bud flush and set, height increment, growth rate, and frost hardiness are extensive as demonstrated by provenance trials and greenhouse experiments (Hurme *et al*., 1997; Andersson & Fedorkov, 2004; Hall *et al*., 2021). Variation in these traits has been correlated with geographic parameters such as latitude or longitude, and with environmental parameters such as temperature, precipitation, and the length of the growing season. Clinal phenotypic variation is a response to spatially varying selection along environmental gradients, which is expected to generate gene frequency clines at associated loci (Savolainen *et al*., 2007). The phenotypical cline, however, can also arise from plastic responses (Vitasse *et al*., 2010; McLean *et al*., 2014), or a combination of both genetic and plastic mechanisms (Corl *et al*., 2018).

**Figure 1:**
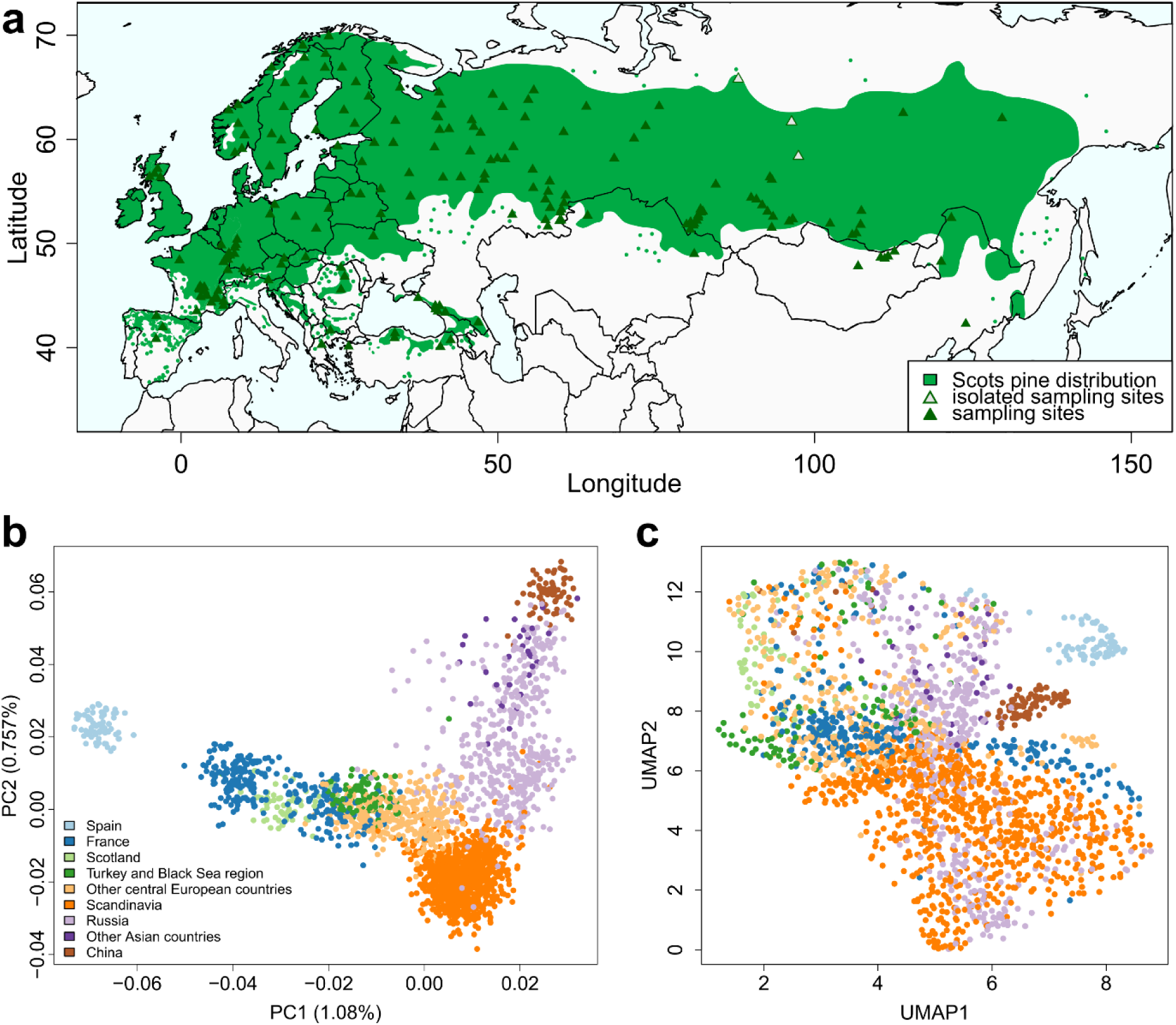
Sampling and genetic distribution in Scots pine. Panel **a**: Geographic distribution of the species with sampling sites. Green triangles represent sampling sites. Three populations indicated by grey triangles were used only in PCA and admixture analyses. Species distribution map from Caudullo *et al*. (2017). Panel **b**: Principal component analysis (PCA) with individuals colored by origin. The same figure colored by country is available in Fig. S4. Panel **c**: Uniform Manifold Approximation and Projection (UMAP) with individuals colored as in panel B.

The current distribution of Scots pine is believed to have been shaped by migration out of several scattered refugia that survived the last glacial maximum (LGM) in southern Europe, Turkey and the Russian Plain (Willis *et al*., 2000; Tóth *et al*., 2017a). Mitochondrial polymorphisms suggest that Scots pine expanded into Fennoscandia from west-central Europe and the Russian Plain 10,000 – 7,000 years ago as they tracked receding ice sheets from ice age refugia (Naydenov *et al*., 2007; Pyhäjärvi *et al*., 2008). However, several recent discoveries question the local extinction of the species during that period and suggest the existence of northern refugia. A tree dated at 11,700 before present (BP) has for example been discovered in central Sweden (Kullman, 2002), which would imply an extremely fast migration rate from the southern Europe refugia if the species had locally disappeared. A vegetation model identified potential *P. sylvestris* refugia during the LGM up to 50°N (Cheddadi *et al*., 2006), based on the European Pollen database and charcoal records showing a continuous presence of conifers during 32,000 - 16,000 BP in central and eastern Europe (Willis & van Andel, 2004).

Rapid colonization from few isolated refugia is expected to leave distinct genetic signature in the extant populations, such as declining genetic diversity and increasing genetic load along migration routes, and genetic lineages that track their recent ancestry (Hewitt, 2000; Excoffier *et al*., 2009). In such a scenario, we would expect lower genetic diversity and higher genetic load in the northern populations of Scots pine and higher differentiation of them from the refugial populations. However, if the species survived in many micro-refugia scattered across the current distribution, we should observe little decline of the diversity in the north, no correlation between genetic load and population expansion, and weak differentiation between populations. Testing these hypotheses requires comprehensive coverage of the species distribution and sufficient genetic information. However, genome-wide patterns of genetic diversity across the species range, as well as the impact of landscapes and environments on population differentiation, remain scarcely studied for Scots pine (but see Sannikov & Petrova, 2012; Tyrmi *et al*., 2020), hindering the recovery of its evolutionary history and genetic signature of local adaptation.

In this study, we conducted genotyping-by-sequencing on 2,321 Scots pine individuals sampled from more than 200 populations across its natural range. With this extensive sampling, we aimed to assess: 1) the composition and distribution of genetic diversity in Scots pine at regional and range-wide scales, and the impact of geography, ecological conditions and gene flow on the spatial patterns, 2) the recent demographic history of the species, and whether expanding populations harbor high genetic load; and 3) the genetic and environmental associations, and the climatic drivers of population differentiation and local adaptation. These analyses enhance our understanding of how diversity correlates with geography, demography, and climate adaptation, and of the adaptive potential of the species in response to current and future environmental changes.

## Material and methods

A more complete methods section is available in Supplementary Methods.

### Sampling and genotyping-by-sequencing (GBS) library preparation

We collected 2895 samples from 211 natural stands covering the entire geographic distribution of *P. sylvestris*. The samples were either needles, buds, or seedlings grown from seeds, collected in native forests to the best of our knowledge. Because the Scots pine genome is large (ca. 23Gbp; Fuchs *et al*., 2008), we used a reduced representation sequencing method called GBS. The library preparation followed the protocol of Pan *et al*. (2015), using a *Pst*I-HF enzyme to digest the DNA. After fragment size selection, paired-end sequencing (2×150 bp) was performed on Illumina HiSeq X Ten (Novogene, UK).

### Genotype likelihoods

After preliminary analyses detailed in Supplementary Methods, we retained 2,321 unrelated individuals from 202 stands (Fig. **1a**, Fig. **S1**, Table S1). Clean reads were mapped on the *P. taeda* genome v.1.01 (Neale *et al*., 2014; Zimin *et al*., 2014) using the Burrows-Wheeler Aligner v. 0.7.17 (Li, 2013). Instead of calling SNPs, we used ANGSD v. 0.935 (Korneliussen *et al*., 2014) to estimate genotype likelihoods, *i.e.* the probability of each genotype at each site and for each individual. We set a minimal individual depth of 5x, a Phred-scaled base quality of 20 and a mapping quality of 40 for an individual site to be considered. We removed the sites with >40% missing data, more than two alleles and an observed heterozygosity higher than 50%.

The samples were sequenced in 13 different libraries, thus slight shift in size selection can cause a certain degree of non-overlap of the sequenced sites between libraries, creating similarity by sequence library instead of geographical origin of the samples. To remove this batch or library effect, we identified the SNPs grouping subsets of samples by library instead of country or region using PCAngsd v. 1.01 (Meisner & Albrechtsen, 2018) and removed them. The final SNP set together with all the invariant sites was used for nucleotide diversity and demographic analyses. We produced another SNP dataset with a minimal allele frequency (MAF) of 0.05 for all the other analyses. We checked that the final SNPs were independent by estimating the linkage disequilibrium between each pair of SNPs using ngsLD v. 1.2.0 (Fox *et al*., 2019).

### Genetic diversity and population structure

To describe the level and structure of genetic diversity at regional and range-wide scales, we first combined the stands having less than six individuals with their closest neighbors to reach at least six individuals per population, the minimal number to obtain reliable diversity estimates (Nazareno *et al*., 2017). Three stands from central Russia that were too distant (>375km; Table S1) to any other sampled stands were left alone. This way, we obtained 123 larger populations, covering the entire distribution (Fig. **1a**, Fig. **S1**). We estimated for each large population the per-site inbreeding coefficient (*F*_IS_), observed (*H*_o_) and expected heterozygosity (*H*_e_) with ANGSD on the MAF 0.05 dataset. We estimated the pairwise nucleotide diversity (*π*) and Tajima’s D statistic (Tajima, 1989) from the folded site frequency spectrum (SFS) based on the full dataset. The 95% confidence interval (CI) of Tajima’s D was estimated by subsampling the data with replacement to produce 100 SFS. The nucleotide diversities at 0-fold sites (*π*_0_, where all mutations would be non-synonymous) and 4-fold sites (*π*_4_, where all mutations would be synonymous) were based on the gene annotations of the *P. taeda* genome.

To characterize the spatial organization of the genetic diversity, we first performed a PCA on the MAF filtered dataset with PCAngsd including all samples. As PCA only performs linear dimension reduction, we added a Uniform Manifold Approximation and Projection (UMAP) analysis which implements non-linear dimension reduction using UMAP v. 0.5.3 (McInnes *et al*., 2018). We then examined the population structure using NGSadmix v. 32 (Skotte *et al*., 2013) with a potential number of clusters *K* ranging from 1 to 20, with 50 replicates for each analysis, and a maximum of 200,000 iterations. We tried to identify the optimal *K* value representing the data using both the Evanno (Evanno *et al*., 2005) and the logarithm method (Pritchard *et al*., 2000).

To describe geographic patterns of genetic differentiation (*F*_ST_) between pairs of large populations, we first estimated *F*_ST_ using ANGSD. We then performed Mantel tests to evaluate the spatial autocorrelation of the results, comparing geographic *vs.* genetic distance between populations (normalized *F*_ST_), genetic *vs.* environmental distance (Mahalanobis distance calculated from the standardized environmental variables), and geographic *vs.* environmental distance using the R package vegan v. 2.5-6 (Oksanen *et al*., 2019) and 9999 permutations.

To account for the impact of isolation by distance (IBD; Wright, 1943) on population differentiation, we examined population structure using tess3r which considers IBD in defining genetic clusters (Caye *et al*., 2016). We ran the tess3r analysis for *K* ranging from 1 to 20, with 20 replicates for each value of *K* and a maximum of 200 iterations.

Finally, to discern spatial patterns of gene flow, we estimated the effective migration surfaces with the python package feems (Marcus *et al*., 2021). This method is based on a stepping-stone model and identifies migration rates between populations that deviate from what would be expected under pure IBD, thereby locating gene flow barriers or corridors. We performed a leave-one-out cross-validation for lambda values (tuning parameter) varying from 10^-6^ to 10^2^, with 20 values tested.

### Demographic history

As historic events can have a significant impact on population genetic diversity and differentiation, we estimated demographic changes of the three most divergent populations in Scots pine (i.e. China, Spain, and northern Norway) using two different approaches. The first, Stairway Plot v. 2 (Liu & Fu, 2015; Liu & Fu, 2020), performs multi-epoch coalescent inference of population size changes through time. It is suitable for demographic analyses where no previous knowledge is available but can only analyze populations independently. The second, fastsimcoal2 v. 2.6 (Excoffier *et al*., 2013; Excoffier *et al*., 2021) implements multi-population demographic models that take divergence and migration among populations into consideration.

We produced folded SFS with ANGSD on the full dataset (including invariant sites) for these three populations and 2D-SFS for each pair of populations. We ran Stairway Plot for each of the three populations separately and for the entire species (2,321 individuals) under the default parameters. We used an average generation time of 20 (Pyhäjärvi *et al*., 2020) and 50 years (Willyard *et al*., 2007; Tóth *et al*., 2019) and a mutation rate of 7 × 10^-10^ mutations per site and year (Willyard *et al*., 2007). We considered the entire allele frequency spectrum with or without singletons.

We tested 20 demographic scenarios with fastsimcoal2 excluding singletons (Fig. **S2**). These 20 models cover most of plausible population history scenarios. Each model was run 50 times, with 100,000 coalescent simulations and 40 expectation-conditional maximization cycles for the calculation of the global maximum-likelihood. The best-fitting model was selected based on the maximum value of likelihood over the 50 independent runs of each model and their Akaike’s weight of evidence (Akaike, 1987). Parameter confidence intervals (95% CI) of the best model were obtained by running 100 parametric bootstraps, with 50 independent runs in each bootstrap. We used a mutation rate of 7 × 10^−10^ mutations per site and year and a generation time of 50 years.

To estimate the potential fitness impact of population expansion (see Results), we first compared the efficacy of purifying selection as measured by *π*_0_/*π*_4_ ratio with the Tajima’s D value for each large population. If demographic expansion is correlated with an increase in genetic load (higher frequency of deleterious mutation), we should observe a negative correlation between the two parameters. We also looked at the gene effects of the SNPs by classifying them using snpEff v. 5.2 (Cingolani *et al*., 2012). We then estimated the additive and recessive genetic load at the individual level as in de Pedro *et al*. (2021) and compared the average values between large populations.

### Genotype and environment association (GEA)

To explore potential signals of genetic adaptation driven by local environmental conditions, we extracted 74 soil, climate, light conditions, and species composition variables from several databases (Table S2) to describe the environment of each of the 123 large populations. Subsequently, we employed a gradient forest (GF) analysis performed with the R-package gradientForest v. 0.1-37 (Ellis *et al*., 2012) to rank them by importance, a random forest concept that evaluates the significance of a predictor for the model. The GF models were validated by a permutation test through 10 different runs using shuffled environmental data. The variables with a correlation coefficient ≥│0.75│ (Spearman’s R) to the top-ranked variable were removed. This was done with every subsequent top ranked variable. The remaining variables were then used as input for the subsequent analyses.

We performed redundancy analyses (RDA; Legendre & Legendre, 2012; Rellstab *et al*., 2015) to evaluate the impacts of three explanatory matrices on allele frequencies across populations. The first matrix was based on the distance between populations through principal coordinates of neighborhood matrix (PCNM) that converts distances to a rectangular representation (Borcard & Legendre, 2002). The second was based on the environmental variables extracted from the GF analysis, and the third was the shared coancestry between populations quantified from the first two UMAP+PCA axis. The first two matrices were first analyzed separately through a forward selection step with the ‘ordistep’ function in the vegan package (Oksanen *et al*., 2019). We ran a set of full and partial RDA models to quantify the impact of each of the three factors. Outliers in RDA were identified by comparing observed and predicted *p*-values distributions (modeled as logistic) of the number of SNPs scored.

We further conducted a BayPass analysis (Gautier, 2015) to identify significant SNPs associated with environmental variables. We first used the core model to establish the covariance of allele frequencies among populations that would be the result of their shared history. We detected outliers that showed strong differentiation among populations based on the observed log-normal distribution of the differentiation X^t^X (*p*-value <0.001, see Fig. **5d**). These outliers are analogous to *F*_ST_ outliers. We then ran the auxiliary covariate model with spatial dependency on the markers using an Ising prior, *b*_is_ = 1, for each of the GF selected environmental variables independently, considering both outliers detected by the X^t^X statistics and other non-differentiated SNPs. We considered a SNP to be correlated with an environmental variable when the standard deviation of the effect estimate did not cross zero (|*β*|-*σ* (*β*) > 0), corresponding to posterior inclusion probability ≥ 0.55. Finally, we identified the environmental variables that showed an association with at least one SNP and ran the previous model with all these variables simultaneously for final result. Significant SNPs identified by any GEA analysis were annotated based on the *P. taeda* genome annotations.

## Results

### Genotype dataset

The 2,321 samples that passed the quality filters averaged 2.42 million (M) mapped reads (0.33 - 49.56 M reads), 2.83 Mbp (million base pairs) with at least 5x depth (1.37 - 7.58 Mbp), and a read depth of 85.26x (17.38 - 953.61x). The final genotype likelihood dataset included 1,432,399 sites, of which 386,426 were SNPs. Filtering for MAF > 0.05 reduced the dataset to 25,830 SNPs. After further checking, we found no evidence of linkage disequilibrium among them. Only 5,623 were in coding sequences, with 2,078 being synonymous mutation, 3,185 being non-synonymous mutation, and 96 being gain or loss of a stop codon.

### Genetic diversity and population structure

We calculated the pairwise nucleotide diversity (π) among samples for each of the 123 large populations and found little variation among them with an average of 0.0062 differences per base pair (Table 1, Fig. **S3**). Across the whole distribution, π values, including π_0,_ π_4_ and the π_0_/π_4_ ratio, decreased slightly with increasing latitude and longitude (Fig. **S4**). The average observed heterozygosity (*H*_o_) was 0.258, and the expected heterozygosity (*H*_e_) was 0.277 (Table 1, Table S1), with a small decrease at high latitudes, but no significant variation along longitude (Fig. **S4**). As anticipated for a highly outcrossing species, we recovered a very low inbreeding coefficient (*F*_IS_) at most locations, averaging 0.059 and ranging from -0.088 to 0.2 (Table 1, Table S1). Slightly elevated *F*_IS_ values were observed in northern Norway (e.g., Alta, Vågå or Kirkesmoen) and in a few isolated populations (Black Sea or Massif Central in France), with a general tendency for marginally higher values at high latitudes (Fig. **S4**). Tajima’s D values were all highly negative (Fig. **S3**, Table 1, Table S1) and significantly different from 0 (one sample t-test; *p*-values < 10^-3^). As these are genome-wide estimations, they can be interpreted as recent population expansion. While no correlation between Tajima’s D with latitude was observed, a weak correlation with longitude was noted, with higher values in Asia (Fig. **S4**).

**Table 1.** Summary of the genetic diversity in Scots pine for each country/region and for the whole distribution. The number of populations (N_pop_) and the number of individuals (N_ind_) for each country or region are given in the two first columns. The overall pairwise nucleotide diversity (π), diversities at the 0-fold (π_0_) and 4-fold (π_4_) sites and Tajima’s D were calculated on the full dataset including invariant sites. The inbreeding coefficient *F*_IS_, the observed and expected heterozygosities (*H*_o_ and *H*_e_) were based on the dataset filtered for MAF 0.05. Six individuals from three very distant Russian populations were not included in these analyses. The values presented are averages of the individual population results, weighted for the number of individuals per population. Tajima’s D significance was tested by a one sample t-test on the 100 replicates and all results were highly significant (*p*-value < 10^-3^). Results by large population are available in Table S1.

| Region / country | $N_{\text{pop}}$ | $N_{\text{ind}}$ | $\pi$ | $\pi_4$ | $\pi_0$ | $H_o$ | $H_e$ | $F_{\text{IS}}$ | Tajima's D |
| --- | --- | --- | --- | --- | --- | --- | --- | --- | --- |
| Whole distribution |  | 2315 | 0.00622 | 0.00890 | 0.00490 | 0.258 | 0.277 | 0.057 | -1.297 |
| Spain | 3 | 70 | 0.00631 | 0.00900 | 0.00504 | 0.265 | 0.273 | 0.027 | -1.369 |
| Scotland | 4 | 59 | 0.00638 | 0.00903 | 0.00504 | 0.284 | 0.292 | 0.020 | -1.032 |
| Western and Central Europe |  | 526 | 0.00650 | 0.00928 | 0.00520 | 0.282 | 0.290 | 0.016 | -1.191 |
| France | 25 | 271 | 0.00655 | 0.00941 | 0.00530 | 0.284 | 0.293 | 0.018 | -1.158 |
| Luxembourg | 1 | 26 | 0.00640 | 0.00913 | 0.00505 | 0.258 | 0.268 | 0.033 | -1.492 |
| Germany | 3 | 37 | 0.00663 | 0.00919 | 0.00519 | 0.278 | 0.287 | 0.019 | -1.306 |
| Austria | 4 | 53 | 0.00643 | 0.00906 | 0.00507 | 0.285 | 0.292 | 0.017 | -1.125 |
| Slovenia | 1 | 17 | 0.00641 | 0.00919 | 0.00502 | 0.278 | 0.280 | 0.007 | -1.252 |
| Slovakia | 1 | 9 | 0.00634 | 0.00900 | 0.00509 | 0.310 | 0.300 | -0.036 | -0.976 |
| Poland | 4 | 68 | 0.00641 | 0.00912 | 0.00513 | 0.266 | 0.277 | 0.026 | -1.325 |
| Belarus | 2 | 20 | 0.00621 | 0.00921 | 0.00495 | 0.299 | 0.296 | -0.018 | -0.892 |
| Estonia | 1 | 6 | 0.00633 | 0.00911 | 0.00506 | 0.346 | 0.325 | -0.072 | -0.776 |
| Bulgaria | 1 | 6 | 0.00632 | 0.00912 | 0.00494 | 0.353 | 0.329 | -0.076 | -0.798 |
| Ukraine | 1 | 13 | 0.00669 | 0.00949 | 0.00534 | 0.270 | 0.288 | 0.045 | -1.401 |
| Southeastern Europe |  | 140 | 0.00644 | 0.00913 | 0.00513 | 0.267 | 0.275 | 0.024 | -1.356 |
| Greece | 1 | 12 | 0.00709 | 0.00982 | 0.00564 | 0.324 | 0.304 | -0.056 | -1.302 |
| Romania | 3 | 60 | 0.00641 | 0.00916 | 0.00518 | 0.261 | 0.274 | 0.039 | -1.377 |
| Turkey | 3 | 68 | 0.00635 | 0.00898 | 0.00500 | 0.263 | 0.271 | 0.026 | -1.347 |
| Eastern Europe and Asia |  | 589 | 0.00610 | 0.00878 | 0.00481 | 0.268 | 0.283 | 0.045 | -1.102 |
| Russia | 34 | 485 | 0.00613 | 0.00885 | 0.00483 | 0.266 | 0.285 | 0.051 | -1.109 |
| Kazakhstan | 2 | 18 | 0.00612 | 0.00892 | 0.00487 | 0.317 | 0.304 | -0.045 | -0.819 |
| Mongolia | 1 | 13 | 0.00616 | 0.00892 | 0.00494 | 0.264 | 0.297 | 0.085 | -1.013 |
| China | 3 | 73 | 0.00591 | 0.00828 | 0.00467 | 0.265 | 0.269 | 0.016 | -1.139 |
| Northern Europe |  | 931 | 0.00610 | 0.00871 | 0.00473 | 0.235 | 0.265 | 0.099 | -1.483 |
| Norway | 9 | 398 | 0.00607 | 0.00862 | 0.00466 | 0.236 | 0.265 | 0.100 | -1.488 |
| Sweden | 10 | 351 | 0.00612 | 0.00880 | 0.00480 | 0.236 | 0.264 | 0.093 | -1.505 |
| Finland | 6 | 182 | 0.00610 | 0.00871 | 0.00472 | 0.233 | 0.267 | 0.109 | -1.428 |

On the PCA performed on the full collection (Fig. **1b**, Fig. **S5a**), we observed a pattern congruent with the geographic distribution: the first axis (PC1) followed a rough South-North direction in west-central Europe, starting from Spain on the left and reaching Scandinavia on the right; the second axis (PC2) followed a more west-east direction for northern European, Russian, and Asian populations. The distribution was rather continuous, with only the Spanish populations being separated from the rest of the samples. The first two PC axes, however, explained less than 2% of the overall genetic variation, meaning that almost all the diversity is shared among all populations. The third and fourth axes, explaining less than 0.4% each, separated the Scottish and Turkish / Black Sea populations from the rest of the samples (Fig. **S5b**). The UMAP analysis confirms the very low level of structure in the data (Fig. **1c**), with only the Spanish and Chinese populations being slightly differentiated from the rest of the sampling, as well as one Romanian population (from Piatra Craiului National Park).

In the clustering analysis, we did not recover a clear optimal number of clusters (*K*): the Evanno method indicated that two groups explained the genetic diversity best, while the logarithm method indicated 20 groups (Fig. **S6**). At *K*=2, we observed a clear gradient between the western (Spanish) and eastern (Chinese) populations, corresponding to the two most distant ones (Fig. **2a**). At *K*=3, the Scandinavian populations formed a new cluster (Fig. **2b**). At *K*=4, a component for western Europe populations appeared. The Russian populations west of the Ural Mountains differentiated from the rest of the Asian populations if we considered five clusters. The Turkish and Black Sea populations split from the western Europe populations at *K*=6 (Fig. **2c**). One Romanian population (from Piatra Craiului National Park) stood out from nine clusters, and the Scottish populations were differentiated when we considered 10 groups. At *K*=12, the Caucasus populations separate from Asia Minor, and no additional pattern emerged at *K* >12 (Fig. **S7**). Similarly, tess3r did not identify a specific number of clusters that best explained the genetic diversity in the population based on cross-validation analysis (Fig. **S8**), and the tess3r results (Fig. **S9**) were very similar to the ones obtained with NGSadmix.

**Figure 2:**
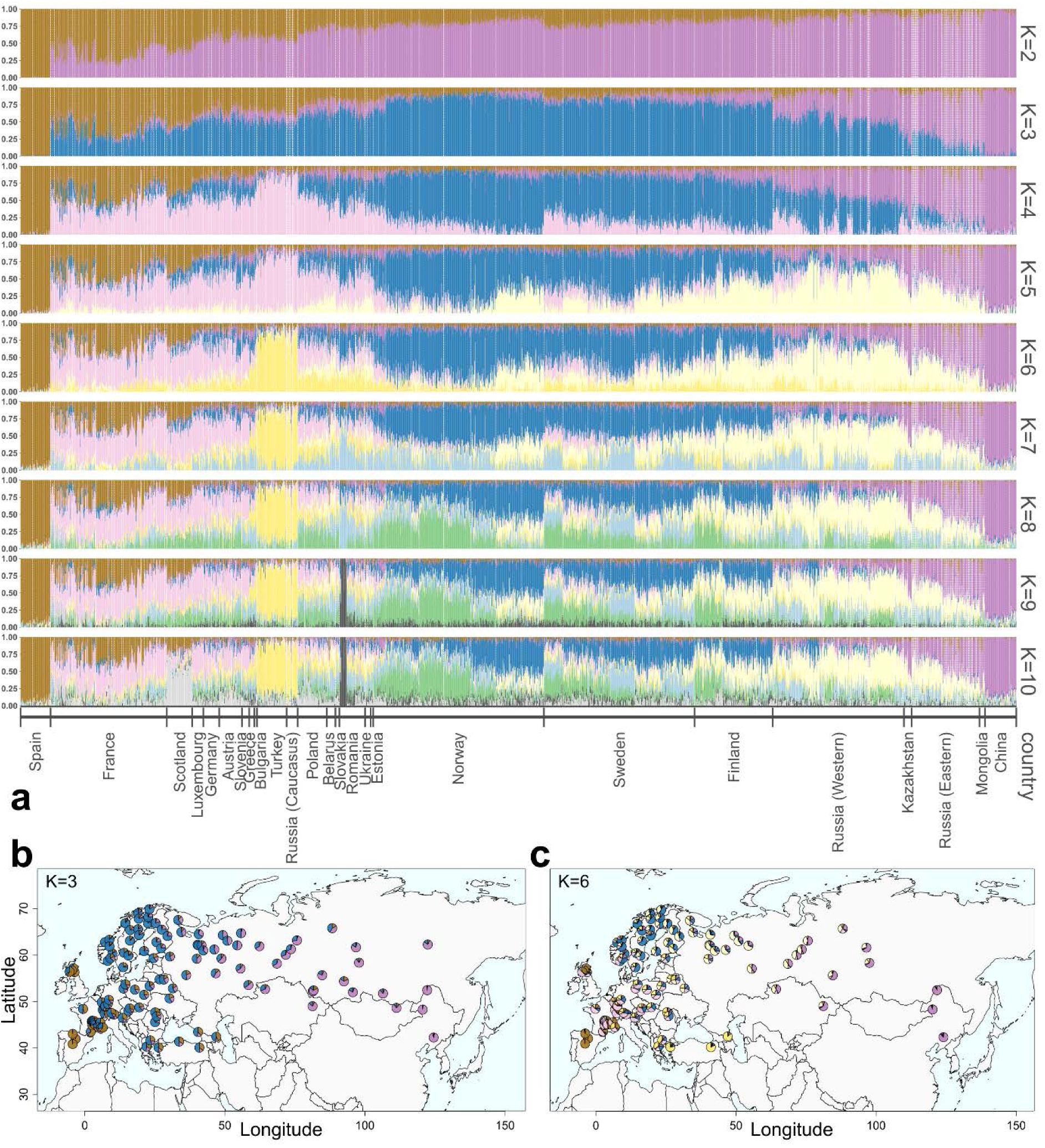
Genetic clustering in Scots pine. Panel **a**: Admixture inference showing the proportion of each cluster component for *K* = 2 - 10. Each color represents one genetic cluster. The populations are ordered by country following the PCA results: from Spain to Scandinavia then to China. In each country, the populations are ordered by longitude except for Scandinavia where they are ordered by latitude. For clustering patterns at *K >*10, see Fig. **S6**. Panel **b** and **c**: Cluster distribution on a map for *K*=3 and *K*=6. The colors follow the result in panel **a**. To reduce the number of pies and increase the readability, the populations represented here are the merged populations (Fig. **S1**).

The pairwise *F*_ST_ between populations ranged from 0.009 to 0.127, and the global *F*_ST_ among the 123 large populations averaged 0.048. In general, the genetic differentiation correlated strongly with the geographic distance separating the populations (Fig. **3a**; Mantel test ρ: 0.586, *p*-value < 2.2×10^-16^), showing a pattern of IBD. By contrast, the correlation of the genetic differentiation with the environmental distance was weak (Fig. **3b**; Mantel test ρ: 0.059, *p*-value: 4.5×10^-7^), indicating no clear pattern of isolation by environment (IBE; Wang & Bradburd, 2014). The correlation between the geographic and environmental distances was also weak (Fig. **3c**; Mantel test ρ: 0.104, *p*-value < 2.2×10^-16^), which means that they can be considered independent in the following analyses. In the feems analysis, we identified several regions with slightly reduced gene flow: the plains around the Pyrenees, Caucasus Mountains and Khingan Mountains, or the North Sea and Aegean Sea (Fig. **3d**).

**Figure 3:**
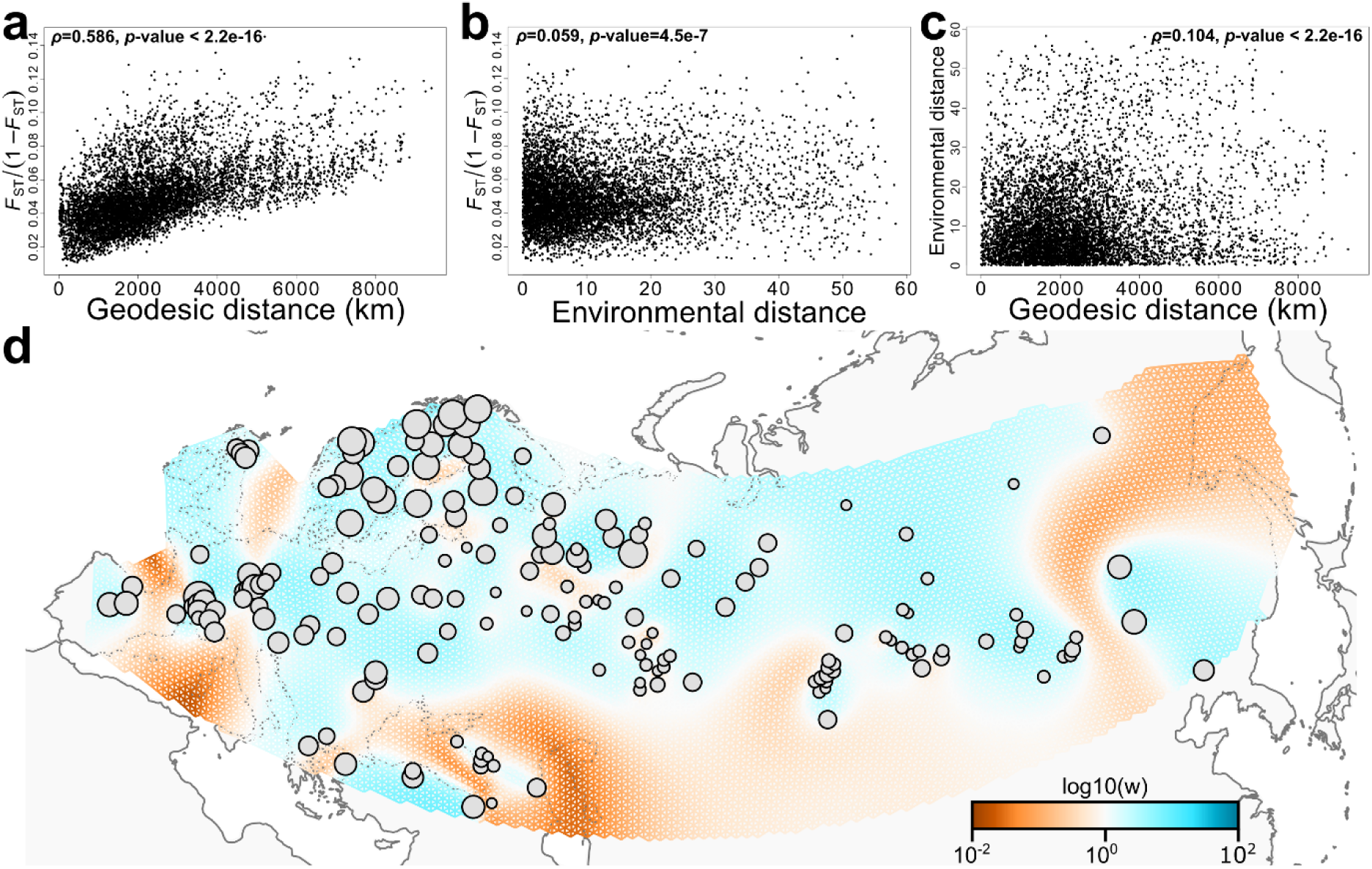
Spatial genetic patterns. Panel **a**: isolation by distance, with Mantel test on geographic *vs* genetic distance between populations. The genetic distance is presented by normalized *F*_ST_ between populations. Panel **b**: isolation by environment with Mantel test on environmental *vs* genetic distance. Panel **c**: Relationship between geographic and environmental distance. Panel **d**: Estimation of the effective migration surface between pairs of populations across the distribution range of Scots pine. Circles correspond to the sampled populations, with radius proportional to the sampling size. Brown areas correspond to a reduction of the gene flow compared to what would be expected given the distance separating populations, and blue areas correspond to an increase of the gene flow.

### Demographic history

When excluding singletons and assuming a generation time of 50 years, we detected with the Stairway Plot 2 analysis an initial phase of population expansion starting around 6 Mya (million years ago, Fig. **4a**, Fig. **S10**). The Spanish population was the first to expand, followed by the Chinese and Scandinavian populations, until they reached their contemporary sizes around 300 kya (thousands of years ago) for the Spanish population, 80 kya for the Chinese population, and 15 kya for the Scandinavian population. Both the Chinese and Scandinavian population sizes peaked at around 150 kya before reaching their contemporary sizes. For the entire species, we observed a sharp increase of the population size starting slightly before 1 Mya and then a rapid contraction at ca. 400 kya. The maximal population size was reached around 300 kya, with a decrease around 20 kya, corresponding to the LGM (Patton *et al*., 2016), followed by a growth reaching 1.1 million individuals around 3,000 years ago. Results for a generation time of 20 years and for analyses with singletons are given in Fig. **S10**.

**Figure 4:**
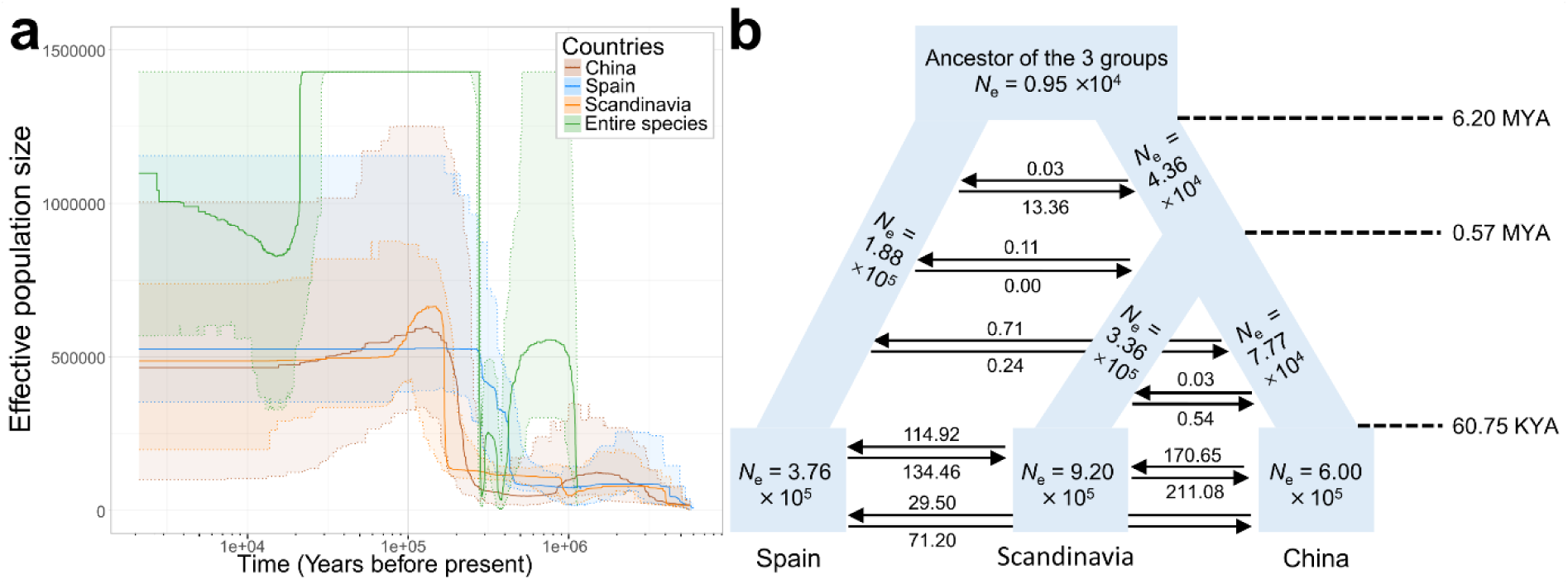
Population history of Scots pine. Panel **a**: Stairway Plot 2 analysis of the three main lineages and of the entire species. The plain lines represent the estimated median effective population sizes (*N*_e_), and the shaded areas correspond to the 95% confidence interval. Panel **b**: fastsimcoal2 analysis. Each block represents a current or ancestral population with their estimated *N*_e_. Arrows indicate the direction of gene flow with the estimated migration rate (number of individuals migrating per generation) labeled above or below the arrow. The timing of the splitting into two groups and their size change are indicated in million years ago (MYA) or thousand years ago (KYA), respectively. The estimates were based on a mutation rate of 7 × 10^−10^ mutations /site /year and a generation time of 50 years, omitting singletons.

Among the 20 demographic models tested with fastsimcoal2, the first two best-fitting models (Model 7 and Model 3) were three-population models with a pattern of isolation-with-migration (Fig. **S2**, Table S3). The expected SFS simulated with these two models reproduced the observed SFS well (Fig. **S11**), suggesting that the recovered demographic parameters were good estimates of the past population history. The estimated demographic parameters from the second best-fitting model (Model 3) overlapped better with those estimated from the Stairway Plot 2 analysis than the first best-fitting model, in terms of *N*_e_ and timing of population size changes (Table S4). This model estimated an ancestral population size *N*_e_ of 9.5 × 10^3^ individuals (95% CI: 9.15 × 10^3^ – 1.48 × 10^4^), which diverged into the Spanish and the ancestral population of the Chinese and Scandinavian groups at 6.20 Mya (95% CI: 5.17 - 6.20 Mya; Fig. **4b**, Table S4). Both populations expanded after the split. The divergence time between the Chinese and Scandinavian populations was estimated at 0.57 Mya (95% CI: 0.49 - 0.63 Mya), followed by expansion. All three populations experienced an instantaneous growth at 60.75 kya (95% CI: 27.95 – 88.07 kya), where *N*_e_ reached 6 × 10^5^, 3.76 × 10^5^, and 9.20 × 10^5^ for the Chinese, Spanish and Scandinavian groups, respectively. The historical gene flow among the three groups was only 0 – 0.71 migrants per generation, while the current gene flow among them was as high as 30 – 211 migrants per generation (Fig. **4b**, Table S4).

The best-fitting model (Model 7, Table S4) uncovered a similar demographic history with estimates of *N*_e_, divergence time, and gene flow among groups in the same orders of magnitude as those of the first model. However, the order of population split was different, as it inferred that the Chinese group split first from the ancestral population. The current *N*_e_ of the Chinese, Spanish and Scandinavian groups were estimated to be 6.60 × 10^5^, 9.97 × 10^5^, and 2.40 × 10^5^, respectively (Table S4), corresponding to a Spanish *N*_e_ twice as high as in the model 3, and a Scandinavian *N*_e_ divided by three.

Contrary to expectation, we did not find any correlation between the π_0_/π_4_ ratio and Tajima’s D, indicating no increase of genetic load in the expanding northern populations (Fig. **S12**). Additionally, neither the recessive nor the additive genetic loads calculated based on gene effects of SNP annotations are correlated with latitude or longitude (Fig. **S13**).

### Genotype and environment association

Based on the GF ranking, we selected the top 21 environmental variables (Table S5) to perform the RDA. Forward selection retained 10 of these variables (Table S5), which were used to produce the environmental distance matrix. For the geographic distance matrix, the forward selection retained 21 of the 64 PCNM-axes. The variance partitioning showed that environment, geographic distance and coancestry together explained 19.6% of the total genetic variance, of which 10.9% was confounded among them (Fig. **5a**). The environmental matrix alone could only explain 0.88% of the total genetic variance. Only the first RDA-axis was identified as significant, and 10 SNPs appeared to be outside of the general clustering and had very low *p*-values (Fig. **5b, c**). However, none deviated from the expected distribution of *p*-values (Fig. **5c**), meaning that these SNPs are not clear outliers. The seven top SNPs appear to have been influenced mainly by arid index, nitrogen content and bio4 (standard deviation of the monthly mean temperatures).

**Figure 5:**
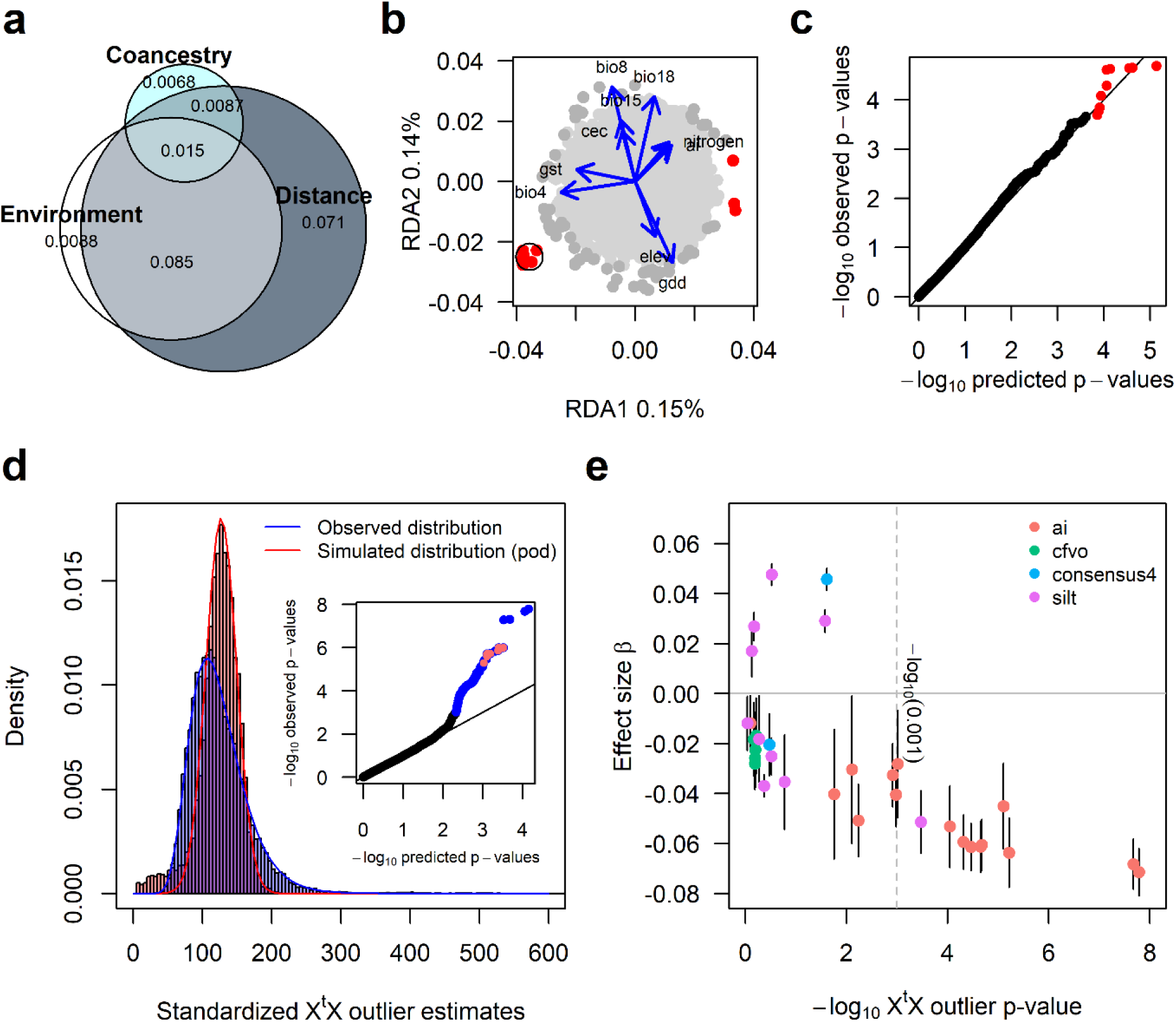
Genotype environment association analyses. Panel **a**: Venn diagram showing the partition of the observed total variance into geographic, co-ancestry and environmental contributions, with the three factors jointly explain 19.5%, while 7.1%, 0.88% and 0.68% independently. Panel **b**: RDA showing the top 10 SNP outliers (●), the circled 7 SNPs were found as X^t^X-outliers in panel **d**. Panel **c**, no p-values of the SNPs from the RDA deviated from the expected distribution. Panel **d**: Comparison of simulated outlier values based on the pseudo-observed data (POD) with observed values. POD estimates are based on the posterior mean and standard deviation of the two parameters of the Beta prior distribution assumed for the (across populations) frequencies of the SNP reference allele of the core model, see material and methods. Inset QQ-plot shows observed vs expected p-values given the log-normal distribution of observed X^t^X outlier estimates (X^t^X values > 311.93). Blue colored dots (● 103) in the inset figures are significant outlier SNPs at p < 0.001 and red highlighted dots correspond to those circled in panel B. Panel **e**: Loci with significant effect size of four climatic/soil variables from the multivariate BayPass analysis. Four variables were found to be associated with two or more loci. Significance was determined when the absolute value of allele effect size (β) minus its standard deviation was larger than zero, |β|-σ (β) > 0, which for this data corresponds to a posterior inclusion probability (PIP) of 0.55 or larger. Because the variables were scaled to a unit’s variance, an allele effect size of -0.06 of ai means that the arid index is 0.06 standard deviations lower when the alternative allele is present. X^t^X outliers are loci that have differentiated allele frequencies across populations.

In the BayPass analysis, we first identified 103 SNPs strongly differentiated among populations (X^t^X outliers), of which seven were also identified in the RDA-analysis (circled in Fig. **5b**). We then detected 33 SNPs (including 11 X^t^X outliers) associated with four environmental variables: aridity index (ai), proportion of silt particles in the fine soil fraction, volumetric fraction of coarse soil fragments (cfvo), and presence of mixed forest (consensus4; Fig. **5e**, Table S6 identified by ^B^). They all deviated from the expected distribution of *p*-values (Fig. **5d**), meaning that these 33 SNPs are strong GEA candidates. Among these 33 SNPs, 17 were in annotated genes whose functions cover synthesis of organic compounds and oligosaccharides, intramembrane protease, splicing and processing of RNA, cell wall reconstruction and stress resistance. However, gene annotations are limited for *P. taeda* and presently absent for *P. sylvestris*, which strongly limits any further interpretation.

## Discussion

### Uniform genetic diversity across Eurasia

Scots pine was able to rapidly expand and (re)establish its distribution across the entire Eurasian continent in the post LGM period. Rapid expansion from a refugial population is expected to leave a spatial pattern characterized by declining genetic diversity and increasing genetic load due to drift and increasing inbreeding on the leading edge of the colonization front. However, our findings in Scots pine deviate from these expectations. Firstly, while there are observable patterns in the diversity (both π and *H*_o_) along latitudes and longitudes, the magnitude of these differences is small. Genetic diversity is reduced by around 10% in northern Europe and Asia compared to the rest of its distribution, indicating a moderate founder effect originating from recolonization. Secondly, the inbreeding coefficient remains close to 0 in most populations, questioning the anticipated increase of inbreeding in geographically marginal populations. Thirdly, there is no discernable signal of genetic load in expanding populations.

The nucleotide diversity is overall high (mean π= 0.0062) compared to other *Pinus* and conifer species (mean π= 0.0013 – 0.0037; Eckert *et al*., 2013; Acosta *et al*., 2019; Shalev *et al*., 2022; Guo *et al*., 2023), or to other boreal tree species with large distribution like the Balsam Poplar, *Populus balsamifera* (mean π= 0.0027; Keller *et al*., 2010), although this difference could partly be due to different sequencing and analysis methods used in each study (Korunes & Samuk, 2021). The average observed heterozygosity (*H*_o_= 0.258) is also high but similar to other Asian pine species (Xia *et al*., 2018; Zhao *et al*., 2020).

The low genetic differentiation in Scots pine is, to our knowledge, unique for a species with comparable distribution ranges. A global *F*_ST_ of 0.048 across its continental distribution uncovered in this study is in agreement with previous results that estimated a *F*_ST_ of 0.02 on average for the western part of Europe (Pyhäjärvi *et al*., 2020; Tyrmi *et al*., 2020; Wachowiak *et al*., 2022a; Milesi *et al*., 2023). We confirm the genetic proximity of the Scottish populations to the mainland ones (Wachowiak *et al*., 2011; Wachowiak *et al*., 2022a), and the presence of genetically distinct populations in Turkey and the Carpathian Mountains detected at high *K* values (Naydenov *et al*., 2007; Dering *et al*., 2017; Tóth *et al*., 2017b). The uniqueness of these populations, and in general the differentiation between European populations, are more visible at the maternally inherited mitochondrial DNA markers due to the more limited dispersal ability of seeds (Tóth *et al*., 2017a). However, gene flow mediated by pollen is sufficiently strong to overcome seed dispersal limitations and to blur the ancestral information in the nuclear genomes of the extant populations.

Associated with the low *F*_ST_, we observed a pattern of IBD, suggesting that spatial distance is a stronger barrier to gene flow than any landforms. Nonetheless, the feems analysis detected barriers to gene flow precisely where we can expect a decrease in pollen exchange: over large bodies of water, or over wide spaces of steppe or broad-leaved woods. The relative isolation of the Spanish and the Caucasian populations identified by feems confirms previous results obtained using other genetic markers (Soranzo *et al*., 2000; Naydenov *et al*., 2007; Semerikov *et al*., 2020; Dering *et al*., 2021; Wachowiak *et al*., 2022b) and can partially be linked to counteracting dominant winds (Kling & Ackerly, 2020). These barriers, even if detectable, are overall faint, and Scots pine can be considered close to panmictic.

This is in stark contrast with Norway spruce that has a distribution largely overlapping the western part of the Scots pine but shows a strong population structure (Chen *et al*., 2019; Milesi *et al*., 2019). Both Norway spruce and Scots pine are wind pollinated with wind dispersed seeds and are mostly outcrossing (Burczyk *et al*., 2004; Robledo-Arnuncio & Gil, 2005; Piotti *et al*., 2009). One reason for their different genetic patterns could be the heavier spruce pollen compared to pine (Di-Giovanni & Kevan, 1991), which reduces the gene flow distance and enhances population genetic differentiation. Other differences in demographic history, number of LGM refugia and ecological amplitude of these species could all impact the distribution of genetic diversity in extant populations resulting in species-specific patterns of genetic structure (Chen *et al*., 2019; Wang *et al*., 2020; Milesi *et al*., 2023).

### Low demographic impact of the LGM

One unexpected result from our demographic analyses was the relatively stable effective population size since 60 kya and little fluctuation during the LGM. The Stairway Plot 2 analysis and the model 3 of the fastsimcoal2 analysis produced similar results regarding the timing of major events and final population sizes. However, the best scenario for the fastsimcoal2 analysis (model 7) inferred a split of the Chinese population before the separation of the Scandinavian and Spanish groups. Although this scenario is plausible, considering the presence of the Scots pine sister species, *P. densiflora,* in the eastern margin of Scots pine range (Mirov, 1967; Saladin *et al*., 2017; Zeb *et al*., 2020; Jin *et al*., 2021), the inferred effective population sizes, with a Spanish *N*_e_ four times larger than the Scandinavian *N*_e_, seem implausible given the lower connectivity of the Spanish populations with the rest of the distribution visible in both the PCA and feems results. We therefore consider that model 3, even if ranked second, is the most plausible based on biological data.

The limited impact of the LGM on the species demography may result from a glacial period too short in duration compared to the evolutionary timescale of the species to exert a significant influence, or from the existence of multiple glacial refugia spread throughout the entire species distribution (including northern latitudes) which could have conserved the entire species genetic variation, or even from the persistence of one large and panmictic population closer to the one observed today. Contrary to the expectation of lower efficacy of purging deleterious mutations in rapidly expending populations, our analyses suggest that the fitness of Scots pine is probably not reduced in the northern expansion limit. This tends to confirm the hypothesis of northern refugia, or even of the persistence of the species in most of its current distribution.

The demographic resilience of Scots pine over the last ice age is not entirely surprising for a cold-resistant species (Hurme *et al*., 1997; Savolainen *et al*., 2004; Hall *et al*., 2021). It faces strong competition from deciduous species when the climate becomes milder (Galiano *et al*., 2010; Boisvert-Marsh & Blois, 2021), making it probably more sensitive to interglacial than to glacial periods. Patches of trees were able to grow on surface debris accumulating on ice (Fickert *et al*., 2007; Zale *et al*., 2018), suggesting that the presence of trees was not limited to ice-free areas during the LGM. If Scots pine had persisted on nunataks or on ice, this would explain the absence of a strong founder effect in our data, as would be expected for a recolonizing species, the mild impact of the LGM detected on the species demography, and the very low differentiation between populations in general. This would also explain the pollen and macrofossil evidence of Scots pines presence at high latitudes very rapidly after and even during the LGM (Huntley & Birks, 1983; Kullman, 2002; Parducci *et al*., 2012; Zale *et al*., 2018). The old image of a few restricted refugia during the LGM is likely inaccurate, and the Quaternary could have been much more forested than initially thought.

The demography of Scots pine recovered in this study supports a strong increase in population size dating back to much older period in Pleistocene or Pliocene (Pyhäjärvi *et al*., 2007; Kujala & Savolainen, 2012; Milesi *et al*., 2023). Similar ancient increases in population size are reported for at least seven other wind pollinated tree species, including three conifers (*Picea abies*, *Pinus pinaster*, *Pinus tabuliformis*) and four angiosperms (*Betula pendula*, *Fagus sylvatica*, *Populus nigra* and *Quercus petraea*) (Xia *et al*., 2018; Milesi *et al*., 2023). Extensive gene flow plays a significant role in shaping these genetic patterns, mitigating the impact of drift and maintaining a large effective population size. Consequently, the success of Scots pine in colonizing new areas after LGM likely arises from a combination of factors, including its resilience, long-distance dispersal ability, and adaptability.

### Signals of GEA

The low genetic differentiation in Scots pine is in stark contrast to the marked phenotypic differences observed among populations, including traits displaying very high *Q*_ST_ (equivalent of *F*_ST_ for phenotypic traits) such as bud set (*Q*_ST_ 0.860; Savolainen *et al*., 2004), growth rate (*Q*_ST_ 0.710; Notivol *et al*., 2007), and cold hardiness (*Q*_ST_ 0.82; Hall *et al*., 2021), demonstrating strong local adaptation (Carlisle & Brown, 1968; Rehfeldt *et al*., 2002). Association studies between markers and traits have reported significant correlations in conifer species (see references in Hall *et al*. (2016)). For example, two GEA studies on Loblolly pine and white spruce, using 20,367 and 6,153 genic SNPs, identified 821 and 307 SNPs linked to environmental variables, respectively (De La Torre *et al*., 2019; Depardieu *et al*., 2021). In contrast, in the case of Scots pine, despite examining 5,623 SNPs in coding sequences, the signals of genetic adaptation to climate gradients were sparse and weak. The overall pattern of isolation by environment barely explained 1% of the genetic differentiation among populations, and only 33 out of 25.8K SNPs displayed a strong association with environmental variables. This paucity of signals may be attributed in part to the limited genome coverage, high level of gene flow among populations, and the lack of IBE observed in our dataset, which collectively impede the recovery of selective signals. In a parallel scenario within the same species, an increase in sequencing to 60,000 genes revealed only one climate-associated SNP, as shown in Tyrmi *et al*. (2020). The weak signals of GEA are theoretically expected for polygenic traits that lack larger allele effects, for which the effect of different co-variance among even minute allele frequency shifts of individual SNPs can lead to large differences in local adaptation (Latta, 1998; Le Corre & Kremer, 2003; Le Corre & Kremer, 2012; Barghi *et al*., 2020). This genetic model implies that a large *Q*_ST_ for a highly polygenic trait can be produced without substantial changes in allele frequencies. It also means that different genotypes potentially can have the same phenotype, thus reducing allele frequency differences among populations and the detection power of frequency-based GEA studies. Previous studies have also shown that despite the lack of significant loci associations, the combined minor effects of many markers can still explain or predict a substantial proportion of the phenotypic variation among genotypes (Hall *et al*., 2021; El-Kassaby *et al*., 2024).

Nonetheless, among the four significant environmental variables that drove the few allele frequency changes detected in this study, the most important one was the aridity index. This is consistent with the ecology of Scots pine, which is known to suffer from drought, resulting in a reduction in tree height in the southern part of the distribution (Hallingbäck *et al*., 2021), shorter and fewer shoots (Taeger *et al*., 2013; Bachofen *et al*., 2021), and increased mortality following severe drought events (Haberstroh *et al*., 2022). Our results highlight the challenges of detecting selection in wild populations of conifer trees. Obtaining a more comprehensive understanding of how natural selection shapes the genetic variation underlying adaptive traits will likely necessitate an approach that considers not only the genetic and environmental aspects but also refines the phenotypic contexts of adaptation under reduced dimensionality of environmental conditions.

## Acknowledgments

We are grateful to the collectors who sent us samples from the field: Philippe Castellar, Sébastien Cecchini (Office National des Forêts), Jean-Paul Collin, Véronique Guérin-Faublée, Caroline Guignier (Monteco), Cédric Hache (Lycée agricole du Haut-Languedoc), Jean-Philippe Hell (Office National des Forêts), Thierry Helminger (Musée national d’histoire naturelle de Luxembourg), Bertrand Jarri (Mayenne Nature Environnement), Isabelle Kerzaon (Université Claude Bernard Lyon 1), Laurent Lathuillière (Office National des Forêts), Marc Oesterle, Benoît Renaux (Conservatoire botanique national du Massif Central), Claude Roulet, Florent Sabatier, Hugues Savay-Guerraz, Hubert Schmuck (Office National des Forêts), Gilles Thébaud (Université Clermont-Auvergne), Hervé Tournier (Parc Naturel Régional du Vercors), Mircea Verghelet and Ionut Bordea (Piatra Craiului National Park), Natalia Demidova (NRIF), Seppo Ruotsalainen (LUKE), Arne Steffenrem and Øyvind Meland Edvardsen (NIBIO). The SAGE data were used with permission of The Center for Sustainability and the Global Environment, Nelson Institute for Environmental Studies, University of Wisconsin-Madison. The computation was enabled by resources provided by the Swedish National Infrastructure for Computing (SNIC) through High-Performance Computing Center North (HPC2N), partially funded by the Swedish Research Council through grant agreement no. 2018-05973. This study was supported by Formas (2018-00842, 2021-02155), Carl Tryggers Stiftelse, T4F program Sweden and a State Contract of the IPAE, Ural Branch of RAS (122021000090-5).

## Author contributions

XRW designed the study and participated in genotyping; JB, WZ and DH conducted genotyping and data analyses; ALC, PA, ADD, OG and HK collected samples from natural populations from their respective regions; AS and VS provided DNA samples; JB, WZ, DH and XRW wrote the manuscript draft, with input from all authors.

## Data availability

The raw read data have been deposited in the NCBI database under the accession number PRJNA976641 (https://www.ncbi.nlm.nih.gov/bioproject/). The correspondence between the file names and the sampling location is available as a supplementary file.

