## Supplementary file for "Scots pine – panmixia and the elusive signal of genetic adaptation"

### Supplementary information

#### Supplementary methods

##### Sampling and genotyping-by-sequencing (GBS) library preparation

We collected 2895 samples from 211 natural forests covering the entire geographic distribution of *P. sylvestris*. The samples were either needles, buds or seedlings grown from seeds. We selected native forests to the best of our knowledge, and the sampled trees in each stand were at least 50 meters apart. Avoiding forests with a recent and documented history of plantation or seeds coming from breeding programs is easy. However, old plantations are almost impossible to trace, as most of them are not documented, without available documentation or available only in the local language. This should nevertheless have little impact on the results given the local origin of the seeds used in the first plantations and the very low structure observed at the continental level (see Results).

We extracted DNA from each individual using an EZNA SP Plant DNA Kit (Omega Biotek). Because the Scots pine genome is large (ca. 23Gbp; Fuchs *et al.*, 2008), we used a reduced representation sequencing method called GBS. The library preparation followed the protocol of Pan *et al.* (2015). Briefly, 200 ng of DNA of each individual was digested with *Pst*I-HF (New England Biolabs) and ligated to sequencing adapters with individual barcodes. Digestion and ligation products from 300 samples (including intra-library and between-library replicates) were pooled into each library, and then purified and PCR amplified. The purified PCR products were separated on a E-gel EX 2% agarose gel (Thermo Fisher Scientific) to recover fragments in the size range of 350 - 450 bp. All libraries were prepared under the same conditions, in the same laboratory. Paired-end sequencing (2x150 bp) was performed on Illumina HiSeq X Ten (Novogene, UK). The source material impacted the individual concentration at the end of the DNA extraction (buds tend to give higher DNA concentration despite a small volume used at the beginning of the extraction) but did not impact the sequencing results: the number of reads were not correlated with the type of material.

##### GBS data filtering

The sequence quality of each library was assessed with FastQC v. 0.11.9 (Andrews, 2010). Reads with > 40% bases having a Phred-scaled quality <20 were discarded by fastp v. 0.23.1 (Chen *et al.*, 2018). Adapter sequences and low-quality bases from the tail of each read were trimmed. Samples in each library were demultiplexed with the process_radtags module of Stacks v.2.0 (Catchen *et al.*, 2013) allowing only one mismatch with the barcode. The first 6 bases of the reads, being the enzyme cutting site, were removed with the fastx_trimmer module of FASTX Toolkit v. 0.0.13 (Gordon & Hannon, 2010). Reads shorter than 41 bp were discarded, corresponding to the five bases of the enzyme recognition site added to the standard minimum of 36 bp. Clean reads were mapped on the *P. taeda* genome v.1.01 (Neale *et al.*, 2014; Zimin *et al.*, 2014) using the Burrows-Wheeler Aligner MEM algorithm v. 0.7.17 with default parameters (Li, 2013). Variants were called using SAMtools and BCFtools v. 1.12 with default parameters (Li *et al.*, 2009; Danecek *et al.*, 2021). Several filtering steps were performed to minimize genotyping errors using BCFtools and VCFtools v. 0.1.16 (Danecek *et al.*, 2011): SNPs located in repetitive regions (reference to *P. taeda* genome v1.01) and with a mapping quality (MQ) <40 were removed; genotypes with a Phred-scaled genotype quality (GQ) <20 or read depth (DP) <5 were masked as missing; loci with a missing rate of >40%, minor allele frequency (MAF) <5%, heterozygosity >50% or not biallelic were also removed. Based on this first SNP dataset, we removed individuals with >60% missing data and individuals with a relatedness coefficient rab higher than 0.5 in each population, calculated using ngsrelate v. 2 (Hanghøj *et al.*, 2019). After these different quality controls, we kept 2,321 individuals from 202 populations (Fig. 1, Table S1).

**Genotype likelihoods**

Instead of calling SNPs, which can bias the results when the sequencing depth is low or when the amount of missing data is high, we used ANGSD v. 0.935 (Korneliussen *et al.*, 2014) to estimate genotype likelihoods for those 2,321 individuals. This allows to keep the probability of each genotype at each site and for each individual in all the subsequent analyses. Because the *P. taeda* genome v.1.01 is highly fragmented, we first concatenated all the regions with at least one mapped read into 20 pseudo scaffolds, separating each region by 300 N. We remapped the reads of each sample against this new reference. We set a maximal depth of 99.5% of the highest depth measured (5022.8x), a minimal depth of 5, a Phred-scaled base quality of 20 and a mapping quality of 40 for an individual site to be considered. We removed the sites with >40% missing data and more than two alleles. We estimated the genotype likelihoods with the GATK method, inferred the major and minor alleles and estimated the allele frequencies. Sites with an observed heterozygosity higher than 50% were discarded.

Because the samples were sequenced in 13 different libraries, we did not have a perfect overlap of the sequenced and missing sites between our samples, and this had an influence on some results (e.g., some samples were grouped by library instead of geographical origin in the principal component analysis; PCA). To remove this so-called batch or library effect, we identified the SNPs grouping subsets of samples by library instead of country or region using PCAngsd v. 1.01 (Meisner & Albrechtsen, 2018) and removed them. We re-ran ANGSD using the final SNP list and all the invariant sites for nucleotide diversity and demographic analyses. We produced another SNP dataset with a minimal allele frequency (MAF) of 0.05 which was used for all the other analyses. We checked that the final SNPs were independent by estimating the linkage disequilibrium between each pair of SNPs using ngsLD v. 1.2.0 (Fox *et al.*, 2019). This software compares the haplotype frequencies of two SNPs and uses a $\chi^2$ test to estimate if they are more similar than expected by chance. We also checked that no SNP deviated from the Hardy-Weinberg expectation using ANGSD. We finally transformed the genotype likelihood dataset to a vcf file for subsequent analyses where genotype likelihoods could not be used.

**Genetic diversity and population structure**

To describe the genetic diversity within and among populations, we first combined the very small populations with their closest neighbors to have at least six individuals per population, the minimal number to obtain reliable diversity estimates (Nazareno *et al.*, 2017), excluding three populations from central Russia that were too distant (>375km; Table S1). This formed 123 large populations, covering as previously the entire distribution (Fig. 1A). We estimated for each large population the per-site observed ($H_o$) and expected heterozygosity ($H_e$) and inbreeding coefficient ($F_{IS}$) with ANGSD on the MAF 0.05 dataset. To estimate the pairwise nucleotide diversity ($\pi$) (Nei & Li, 1979) and Tajima's D statistic (Tajima, 1989), we generated the folded site frequency spectrum (SFS) from the full dataset (including invariant sites) with ANGSD and realSFS, and estimated the nucleotide diversity with the realSFS saf2theta command and the thetaStat program in ANGSD. To obtain the 95% confidence interval around the Tajima's D value, we estimated 100 SFS by subsampling the data with replacement. Nucleotide diversity at 0-fold ($\pi_0$) and 4-fold ($\pi_4$) sites (genes nucleotides whose mutations are never or always synonymous, respectively) were defined based on the gene annotations on the original reference.

To characterize the spatial organization of genetic diversity, we first performed a PCA on the MAF filtered dataset with PCAngsd including all samples. As PCA only performs linear dimension reduction, we added a Uniform Manifold Approximation and Projection (UMAP) analysis which implements non-linear dimension reduction using UMAP v. 0.5.3 (McInnes *et al.*, 2018). We used a local neighborhood value of 15, and a minimal distance of 5 and ran the analysis with and without the PCA coordinates of each sample on all the PC axes. The results were similar and only the results without PCA data are presented. We then examined the population structure using NGSadmix v. 32 (Skotte *et al.*, 2013) with a potential number of clusters $K$ ranging from 1 to 20, 50 replicates for each analysis, and a maximum of 200,000 iterations. The results were combined with the R package pophelper v. 2.3.0 (Francis, 2017) in R v. 4.1.0 (R Core Team, 2021). We tried to identify the optimal $K$ value representing the data using both the Evanno and the logarithm methods in CLUMPAK (Pritchard *et al.*, 2000; Evanno *et al.*, 2005; Kopelman *et al.*, 2015).

To describe geographic patterns of genetic differentiation ($F_{ST}$) between pairs of large populations, we first estimated $F_{ST}$ using ANGSD. To evaluate the spatial autocorrelation of the results, we performed a Mantel test on genetic distance between populations (as normalized population differentiation $F_{ST}$) *vs*. geographic distances using the R package vegan v. 2.5-6 (Oksanen *et al.,* 2019) in 9999 permutations. $F_{ST}$ values between populations were obtained from ANGSD and the geographic distances from the geosphere package v. 1.5-10 in R (Hijmans, 2019). Similarly, we performed Mantel tests on genetic *vs* environmental distance, and geographic *vs* environmental distance. Environmental distances between populations were obtained from RDA, see next section.

To account for the impact of isolation by distance (IBD) on population differentiation, we examined population structure using tess3r which considers IBD in defining genetic clusters (Caye *et al.*, 2016). We ran the tess3r analysis for $K$ ranging from 1 to 20, with 20 replicates for each value of $K$ and a maximum of 200 iterations. We estimated the root mean-squared errors for cross-validation.

Finally, to identify spatial gene flow barriers across the distribution range, we estimated the effective migration surfaces with the python package feems (Marcus *et al.*, 2021). Effective migration surfaces are an estimate of migration rates along graph edges covering the studied area that would produce genetic dissimilarities similar to those observed in the data using an idealized stepping-stone model. We transformed the vcf into a bed file with PLINK v. 1.9b5 (Chang *et al.*, 2015; Purcell & Chang, 2017) and used a triangular grid covering the species distribution at a resolution 7 created with the package dggridR v. 3.0.0 in R (Barnes & Sahr, 2021), corresponding to triangles of 1,557 $km^2$. We performed a leave-one-out cross-validation for lambda values (tuning parameter) varying from $10^{-6}$ to $10^2$, with 20 values tested.

**Demographic history**

To understand how Scots pine colonized its current distribution and how this could have impacted the current genetic diversity and differentiation, we inferred past demographic changes using two complementary approaches. The first, Stairway Plot 2 (Liu & Fu, 2020), performs multi-epoch coalescent inference of population size changes through time. It is suitable for demographic analyses where no previous knowledge is available but can only analyze populations independently. The second, fastsimcoal2 (Excoffier *et al.*, 2013; Excoffier *et al.*, 2021), implements multi-population demographic models that take divergence and migration among populations into consideration.

Following the PCA and admixture results, we estimated the demography of the three most divergent populations in Scots pine (i.e. China, 73 individuals; Spain, 70 individuals; and northern Norway, 168 individuals coming from the Beiarn, Hemne and Molde populations). We produced folded SFS by ANGSD on the full dataset (including invariant sites) for these three groups, which included 1,431,145; 1,431,110 and 1,429,721 sites for China, Spain and Norway, respectively. We also produced 2D-SFS which included 1,430,535 sites for the China-Spain pair, 1,429,276 sites for the Norway-Spain comparison, and 1,429,401 sites for the Norway-China pair.

We ran Stairway Plot for each of the three populations separately and for the entire species (2,321 individuals) under the default parameters and tested four different numbers of random breakpoints using 200 replicates. We used an average generation time of 20 (Pyhäjärvi *et al.*, 2020) and 50 years (Willyard *et al.*, 2007; Tóth *et al.*, 2019) and a mutation rate of $7x10^{-10}$ mutations per site and year (Willyard *et al.*, 2007), which is an intermediate value among the different estimates produced so far (Pyhäjärvi *et al.*, 2007; Willyard *et al.*, 2007; Buschiazzo *et al.*, 2012; De La Torre *et al.*, 2017). Because Stairway Plot analyses can be sensitive to rare variants which could potentially be due to sequencing/genotyping errors (Liu & Fu, 2020), we considered the entire allele frequency spectrum with or without singletons.

We tested 20 scenarios with fastsimcoal2, all of which began with the ancestral population splitting into three different populations but differed in terms of: (i) whether the split events occurred simultaneously or not; (ii) whether divergence occurred with migration or not; (iii) levels of gene flow between two populations, (iv) the mode of population size change after the split, and (v) whether admixture was involved (Fig. S2). Considering the low confidence of rare alleles, we ran fastsimcoal2 based on SFS with "--nosingleton". Each model was run 50 independent times, with 100,000 coalescent simulations as well as 40 expectation-conditional maximization cycles for the calculation of the global maximum-likelihood. The best-fitting model was selected based on the maximum value of likelihood over the 50 independent runs of each model and their Akaike's weight of evidence (Akaike, 1987). The goodness of fit of the best model was tested by comparing the observed SFS with the expected SFS, which was obtained by 100,000 coalescent simulations under the maximum-likelihood estimates of population parameters. Parameter confidence intervals (95% CI) of the best model were obtained by running 100 parametric bootstraps, with 50 independent runs in each bootstrap. We used a mutation rate of $7 \times 10^{-10}$ mutations per site and year and a generation time of 50 years to convert model parameters to absolute values.

To estimate the potential fitness impact of the population expansion detected in these two analyses (see Results), we first compared the $\pi_0/\pi_4$ ratio with the Tajima's D value for each large population (Rougemont *et al.*, 2020). If demographic expansion is correlated with an increase in the genetic load (higher frequency of deleterious mutation), we should observe a negative correlation between the two parameters. We also looked at the impact of the SNPs on the genome by classifying them using snpEff v. 5.2 (Cingolani *et al.*, 2012). We then estimated the additive and recessive genetic load at the individual level as in de Pedro *et al.* (2021) and compared the average values between large populations.

**Genotype and environment association (GEA)**

With its large distribution covering wide environmental gradients, we anticipate the presence of genetic adaptation in *P. sylvestris*. To identify important environmental variables responsible for the overall genomic pattern, we first performed a gradient forest analysis with the R-package gradientForest (GF; Ellis *et al.*, 2012). We then used two different methods, RDA (Forester *et al.*, 2018) and BayPass (Gautier, 2015), to test whether the sampled genetic variation correlated with biotic and abiotic variables. A GF analysis develops non-parametric functions that describe allele frequency turnovers along environmental gradients. We extracted 74 soil, climate, light conditions, and species composition variables from several databases (Table S2) describing the environment of each of the large populations (123) and used a GF analysis to rank them by importance, a random forest concept describing how valuable a predictor is for the model. The GF models were validated by a permutation test through 10 different runs using shuffled environmental data. The variables with a correlation coefficient $\geq |0.75|$ (Spearman's R) to the top-ranked variable were removed. This was done with every subsequent top ranked variable. The remaining variables were then used as input for the RDA and BayPass analyses.

RDA is a method that tries to find the best combination of environmental variables (among a constrained list) that can explain population allele frequencies, assuming that the environmental variables are independent, and that the allelic distribution is dependent on those variables. We built three explanatory matrices to test their influence on allele frequencies across populations. The first matrix was based on the distance between populations through principal coordinates of neighborhood matrix (PCNM) that converts distances to a rectangular representation (Borcard & Legendre, 2002). The first step converts coordinates into geodesic distances in kilometers through the 'geoXY' R-function in the SoDA package in R (Chambers, 2013). Then all pairwise Euclidean distances between each population is calculated ('dist' R-function), which in turn is converted to a PCNM through the 'pcnm' function in the vegan package The second matrix was based on the environmental variables extracted from the GF analysis. The third matrix was based on the first two PCA + UMAP axis to quantify the signal from coancestry. The first two matrices were first analyzed separately through a forward selection step where the allele frequencies were set as response variables with the 'ordistep' function in the vegan package. In these steps we want to maximize the variance explained by a set of predictors. For each step, the predictor that has the greatest explanatory power of the observed variance was added and the model was evaluated. We ran RDA on the distance and environment models and a partial RDA-model where the environmental factors are conditional on the geographic distance between populations and the shared coancestry. This allowed us to estimate the environmental factors that drive allele frequency change while accounting for the spatial and migratory relationship. Outliers in RDA were detected by comparing observed and predicted *p*-values distributions (modeled as logistic) based on the number of SNPs scored.

BayPass builds on previous methods established in BayEnv2 (Günther & Coop, 2013) and provides more robust estimates of population covariance matrixes. It includes a calibration method for the outlier ($X^tX$) statistic and uses the linkage disequilibrium information to better pinpoint loci in significant associations with environmental co-variables. We first used the core model to establish the covariance of allele frequencies among populations that is a result of their shared history. We ran the model with 200 000 iterations and a thinning rate of 20 (-nval 10 000 and -thin 20) with the remaining parameters set to default values (burnin 5000, max number of pilot runs 20 and pilot run length 500). The core model was run three times to ensure stable results. We then created a pseudo-observed dataset (POD) based on the posterior means of the $a_\pi$ and $b_\pi$ parameters of the beta prior distribution assumed for SNP reference allele frequency across populations, $\pi_i$, produced by the core model.

$$\pi_i | a_\pi, b_\pi \sim \beta(a_\pi; b_\pi)$$

This is then used to simulate a reference set of $X^tX$ differentiation values (Günther & Coop, 2013) that can be used to calibrate observed $X^tX$ values. We compared the observed and POD distributions of $X^tX$-values but due to the very different types of distributions we relied on the observed log-normal distribution of $X^tX$-values to establish significance threshold of outliers (*p*-value <0.001, see Fig. 5D). These outliers are analogous to $F_{ST}$ outliers. We then ran the auxiliary covariate model with spatial dependency on the markers using an Ising prior, $b_{is} = 1$, with the same MCMC sampling parameters as in the core model above, for each of the GF selected environmental variables independently, considering both outliers detected by the $X^tX$ statistics and other SNPs not differentiated between populations. We considered that a SNP was correlated with an environmental variable when the standard deviation of the effect estimate did not cross zero ($|\beta|-\sigma(\beta) > 0$), which corresponds to posterior inclusion probability ≥ 0.55. Finally, we identified the environmental variables that showed an association with at least one SNP and ran the previous model with all these variables simultaneously for the final result. SNPs identified by any GEA analysis were annotated based on the *P. taeda* genome annotations.

#### Supplementary figures

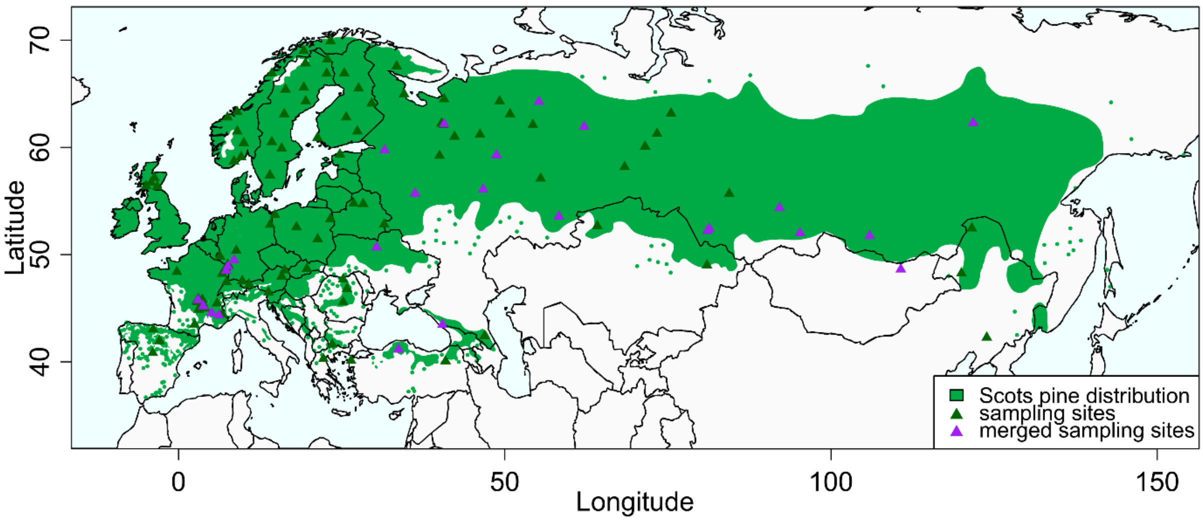


**Figure S1**: *Pinus sylvestris* distribution and merged sampling sites. Green triangles represent sampling sites whose individuals were used in all analyses. Purple triangles represent populations used for genetic diversity description (pairwise genetic differentiation, heterozygosity, fixation index…) and genotype-environment association. Species distribution from Caudullo *et al.* (2017). The map was created using the R packages rgdal v. 1.5-28 (Bivand *et al.*, 2021), mapplots v. 1.5.1 (Gerritsen, 2018) and mapproj v. 1.2.8 (McIlroy, 2022).

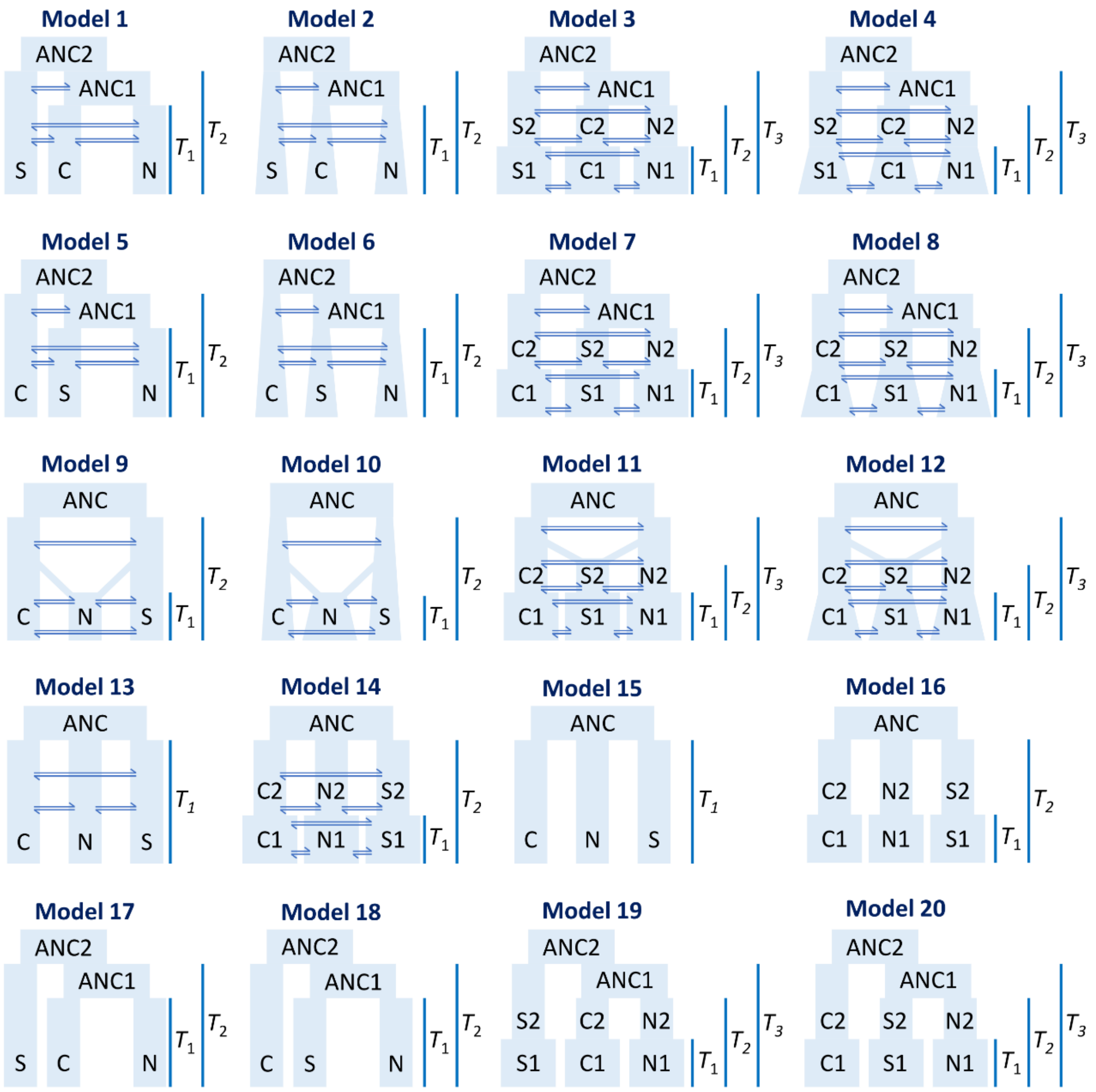


**Figure S2**: Demographic models used in model selection in the fastsimcoal2 analysis. Models 1–14: isolation with gene flow; Models 15–20: isolation without gene flow. Models 1–4, 17 and 19: the Spanish group (S) diverged first; Models 5–8, 18 and 20: the Chinese group (C) diverged first; Models 9–12: the Spanish and Chinese group diverged first, the Scandinavian group (N) was the product of admixture between them; Models 13–16: the Scandinavian, Spanish and Chinese groups diverged at the same time. In model 1, 5, 9, 13, 15, 17 and 18, all groups have constant population sizes after splitting. Models 2, 6 and 10: all groups experienced exponential population size change after splitting. Models 3, 7, 11, 14, 16, 19 and 20: each group experienced an instantaneous population size change at a recent time, after the splitting of the different populations. Models 4, 8 and 12: all groups expanded exponentially at a recent time after the splitting of the different populations.

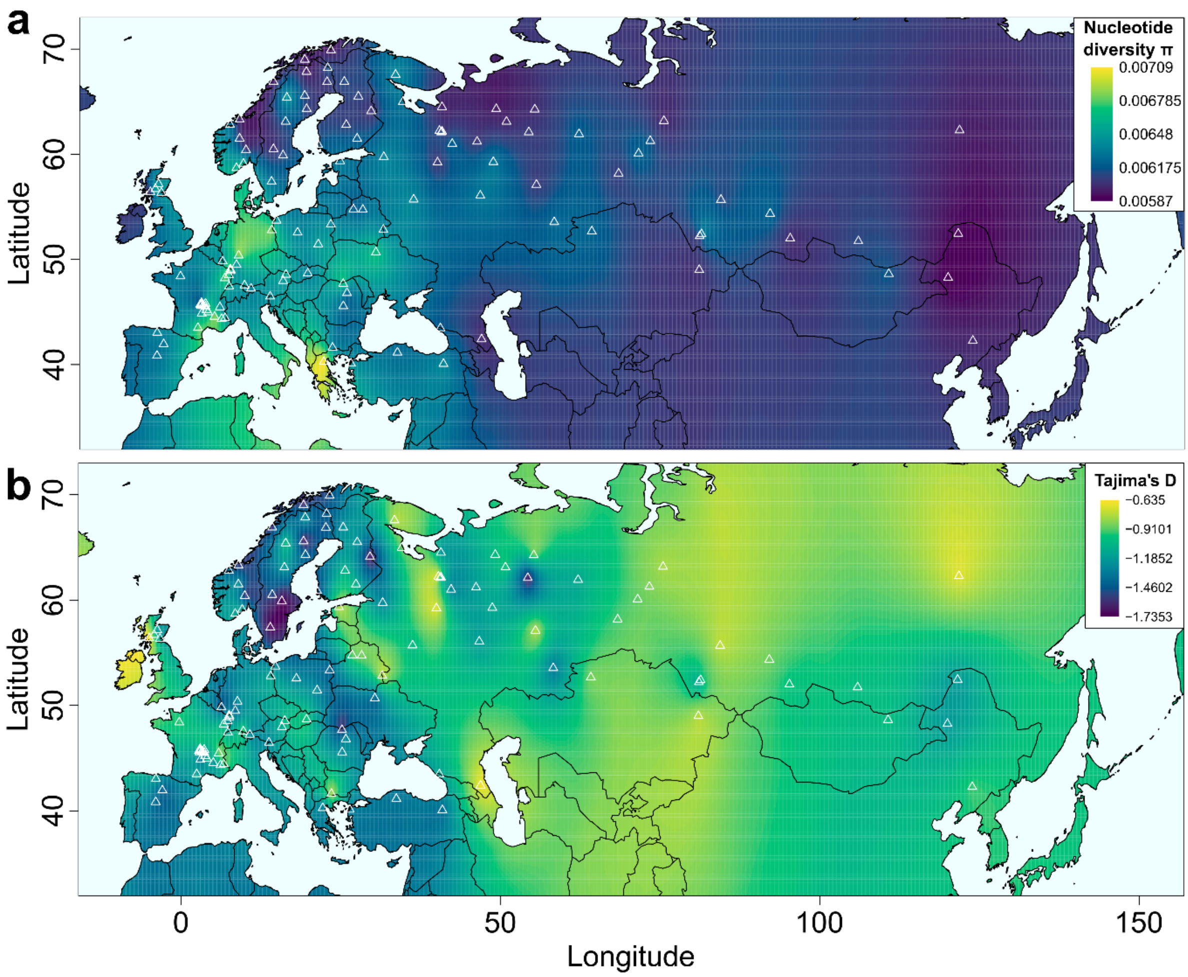


**Figure S3**: Distribution of nucleotide diversity (**a**) and Tajima's D (**b**) values based on estimations for each of the 123 large populations (white triangles) and interpolated using the kriging package v. 1.2 in R (Olmedo, 2022).

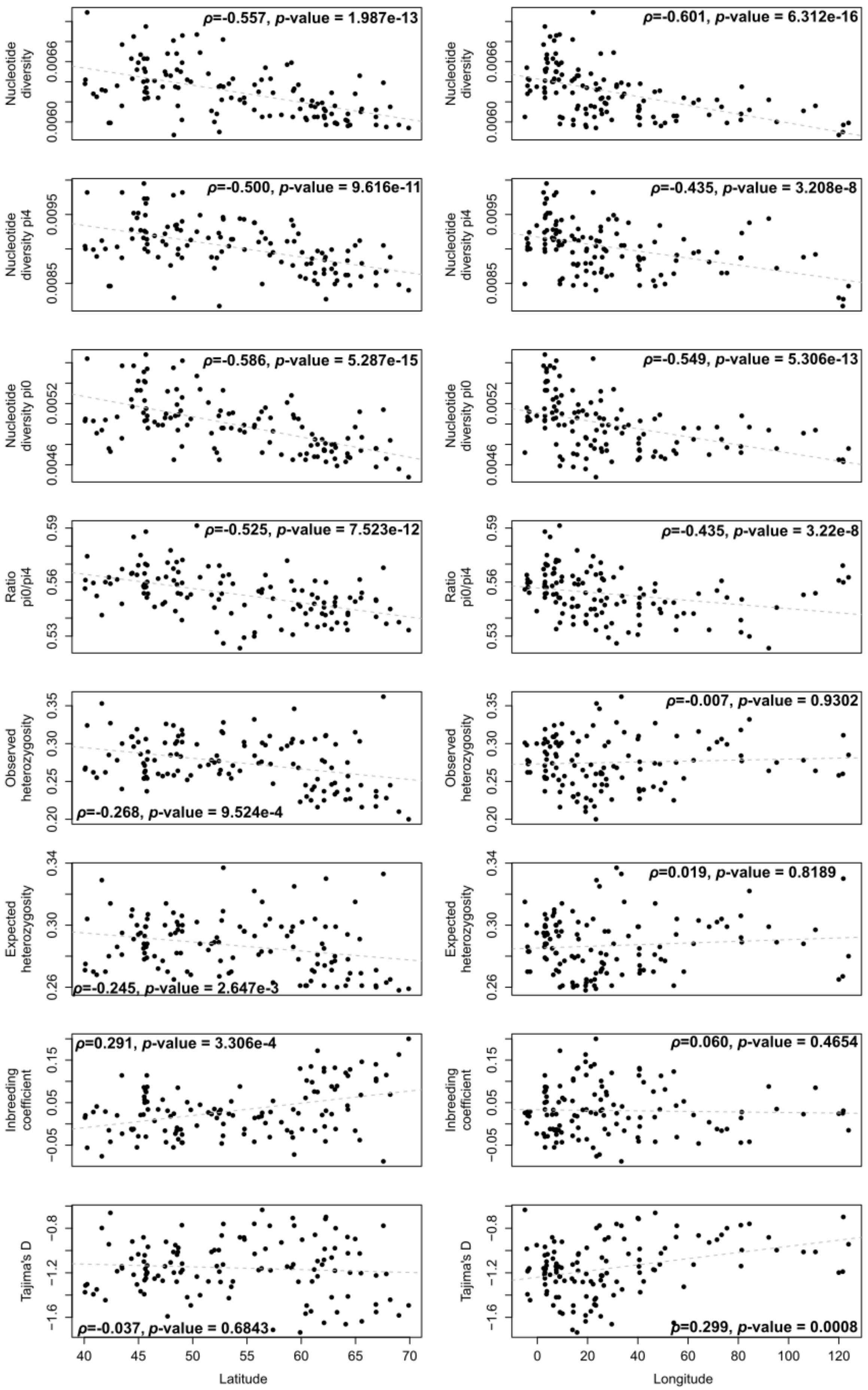

ρ=-0.557, p-value = 1.987e-13
ρ=-0.601, p-value = 6.312e-16
Nucleotide diversity
ρ=-0.500, p-value = 9.616e-11
ρ=-0.435, p-value = 3.208e-8
Nucleotide diversity pi4
ρ=-0.586, p-value = 5.287e-15
ρ=-0.549, p-value = 5.306e-13
Nucleotide diversity pi0
ρ=-0.525, p-value = 7.523e-12
ρ=-0.435, p-value = 3.22e-8
Ratio pi0/pi4
ρ=-0.268, p-value = 9.524e-4
ρ=-0.007, p-value = 0.9302
Observed heterozygosity
ρ=-0.245, p-value = 2.647e-3
ρ=0.019, p-value = 0.8189
Expected heterozygosity
ρ=0.291, p-value = 3.306e-4
ρ=0.060, p-value = 0.4654
Inbreeding coefficient
ρ=-0.037, p-value = 0.6843
ρ=0.299, p-value = 0.0008
Tajima's D
Latitude
Longitude

**Figure S4**: Nucleotide diversity at all sites, 0-fold and 4-fold sites, ratio between nucleotide diversities at 0-fold and 4-fold sites, observed and expected heterozygosity, inbreeding coefficient and Tajima's D for each population, in relation to its latitude (on the left) and longitude (on the right). The Spearman correlation coefficients and the *p*-values are indicated for each figure. The regression lines (grey doted lines) are presented for illustration only, as the residuals of the linear model did not follow a normal distribution with homoscedasticity.

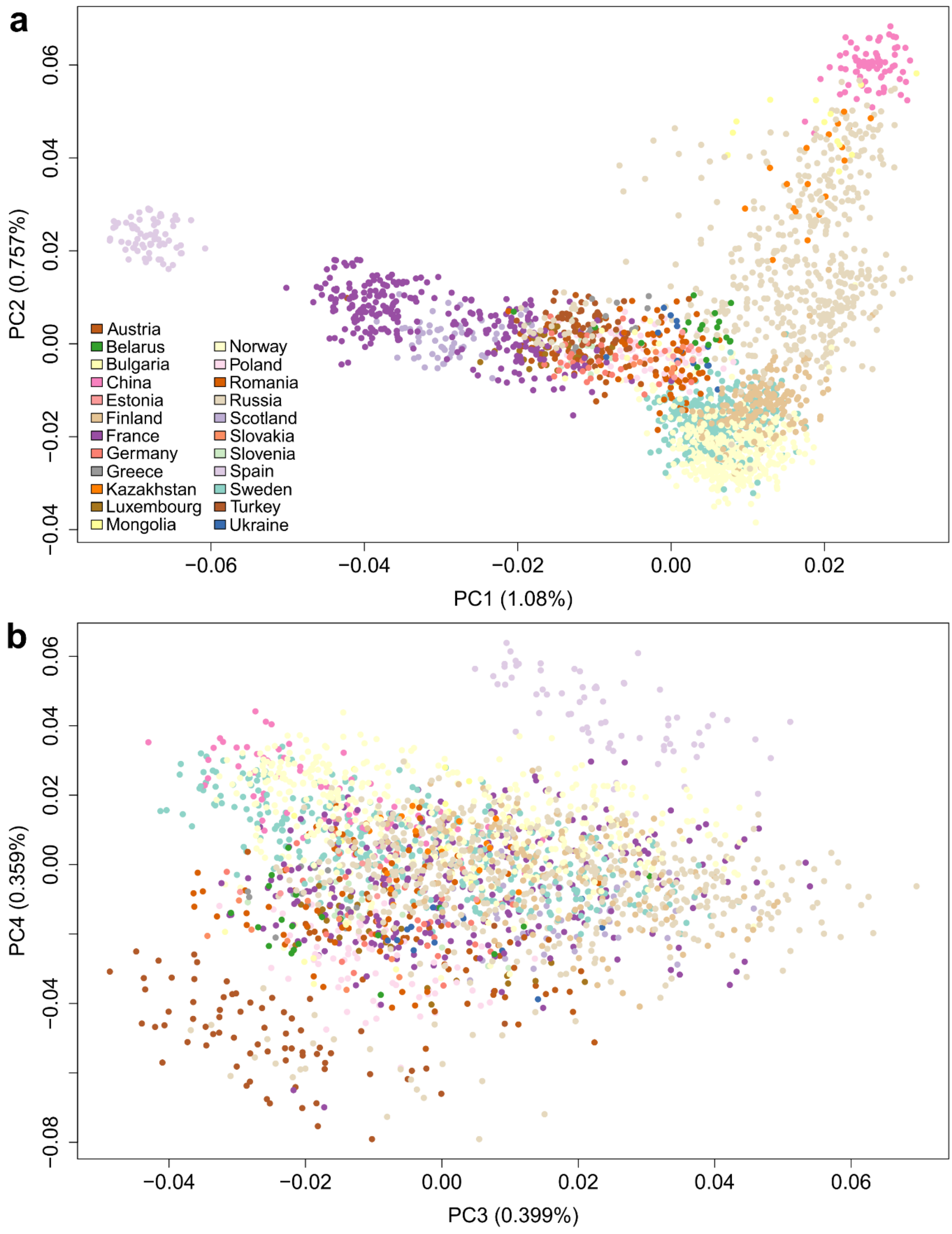


**Figure S5**: Principal component analysis of the 2321 *Pinus sylvestris* individuals colored by country of origin. The percentage of variance explained by each axis is indicated between brackets. Panel **a**: PC1 and PC2. Panel **b**: PC3 and PC4.

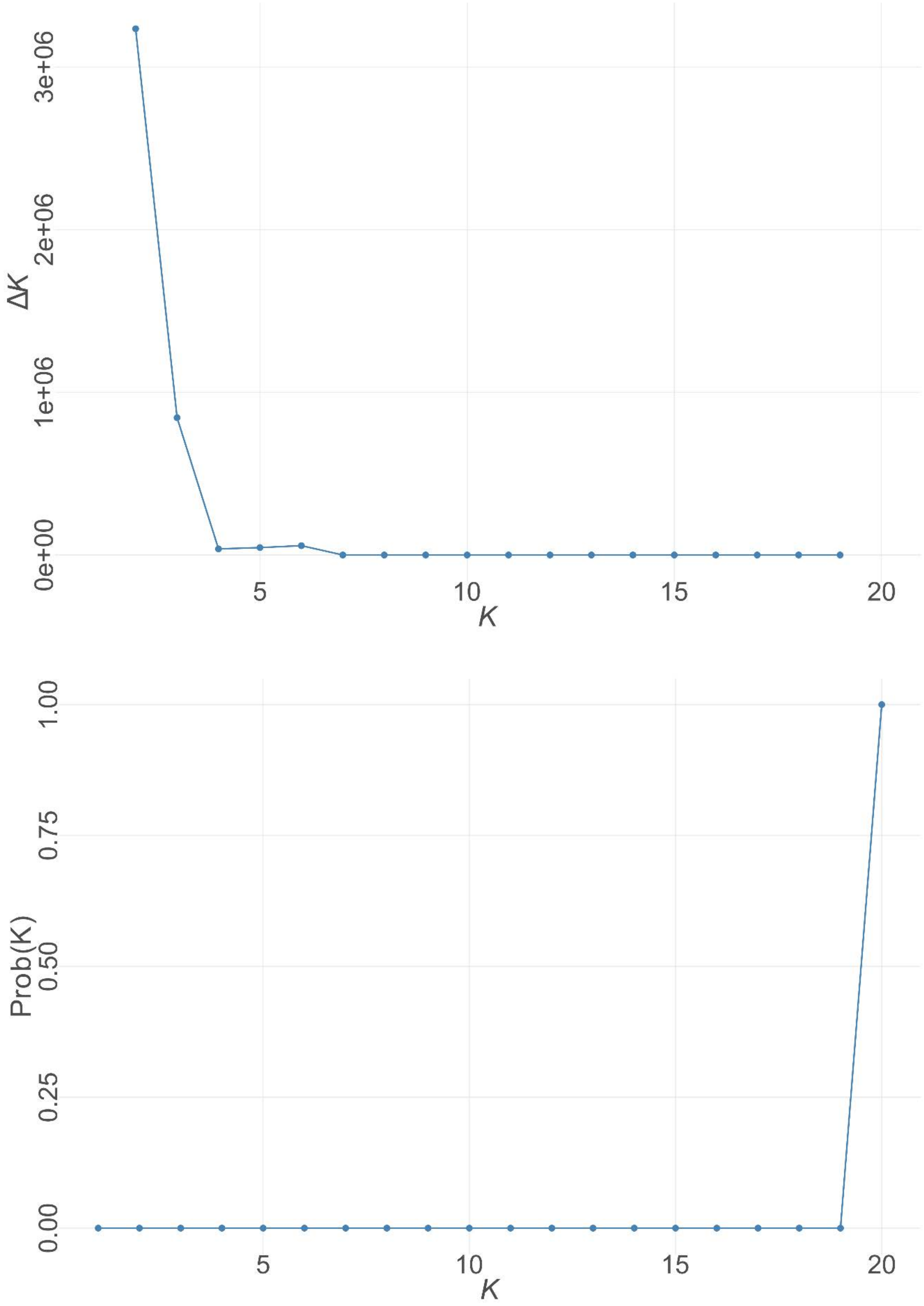


**Figure S6**: Estimation of the best *K* value for the ngsAdmix analysis, using the Evanno method (upper) or the Pritchard method (lower).

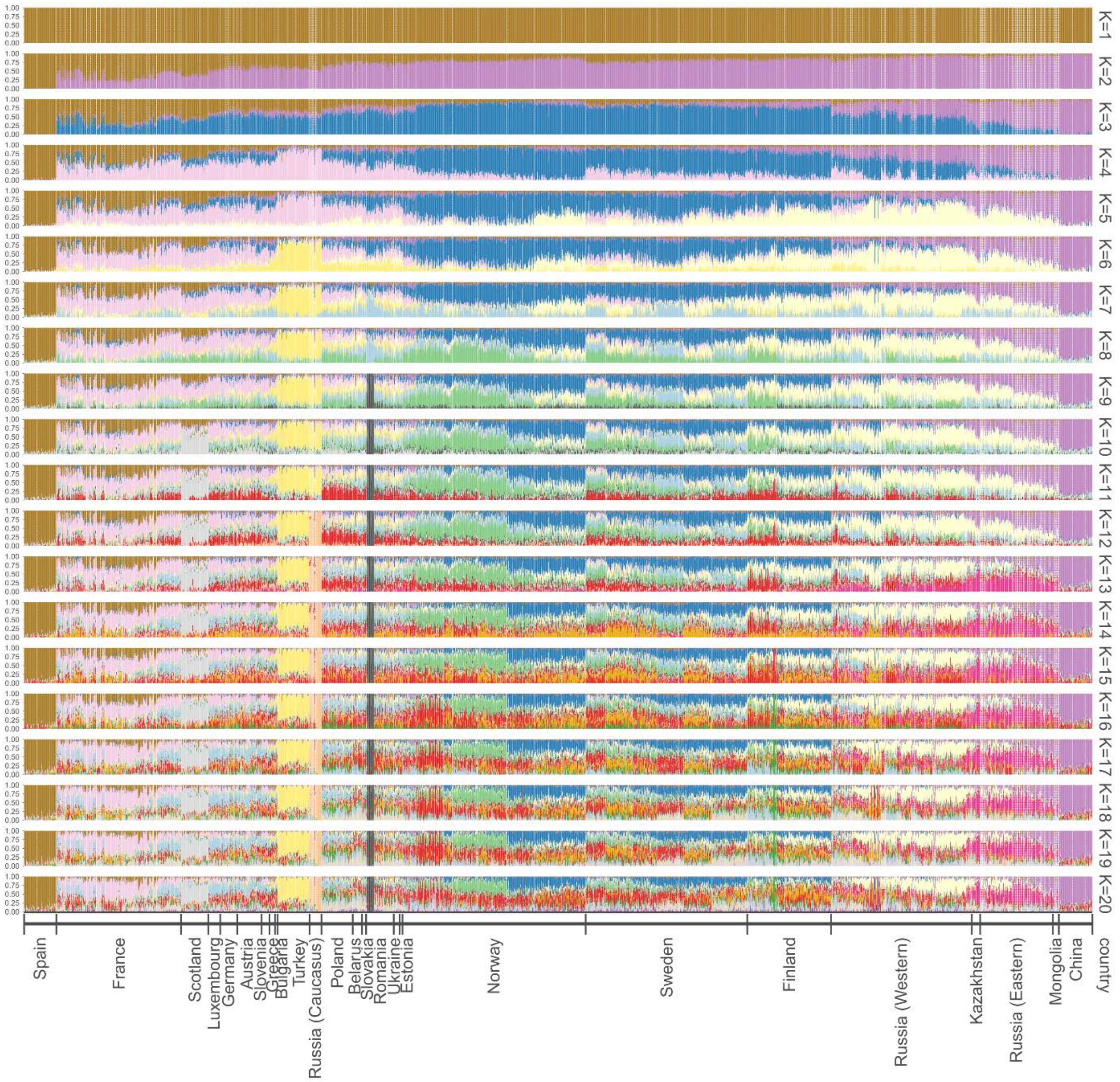


**Figure S7**: Admixture inference showing the proportion of each cluster component for *K* varying between 1 and 20 groups. The populations are ordered by country following the PCA results: from Spain to Scandinavia then to China. In each country, the populations are ordered by longitude except for Scandinavia where they are ordered by latitude.

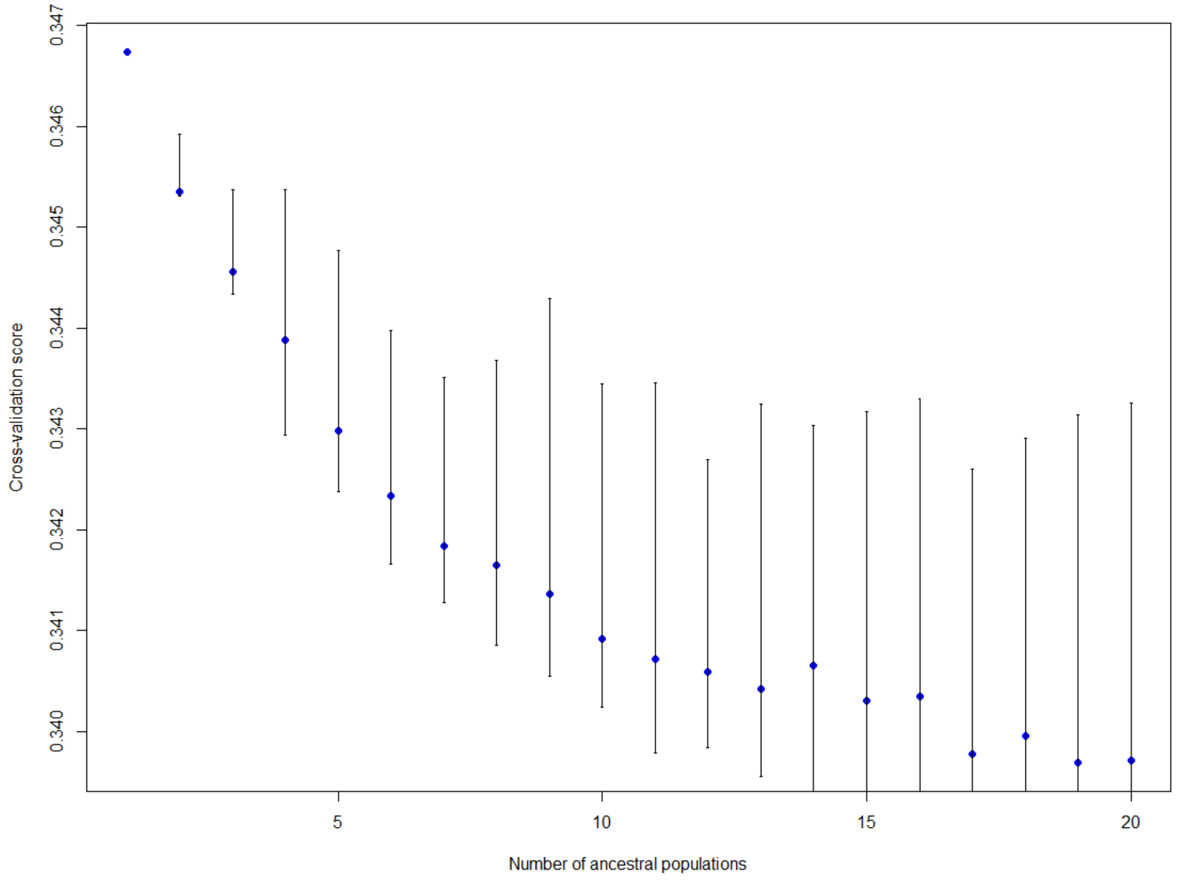


**Figure S8**: Root mean-squared errors calculated during the tess3 analysis for a number of populations ranging from one to 20. Error bars corresponding to the variance observed across the 50 different replicates of the analysis.

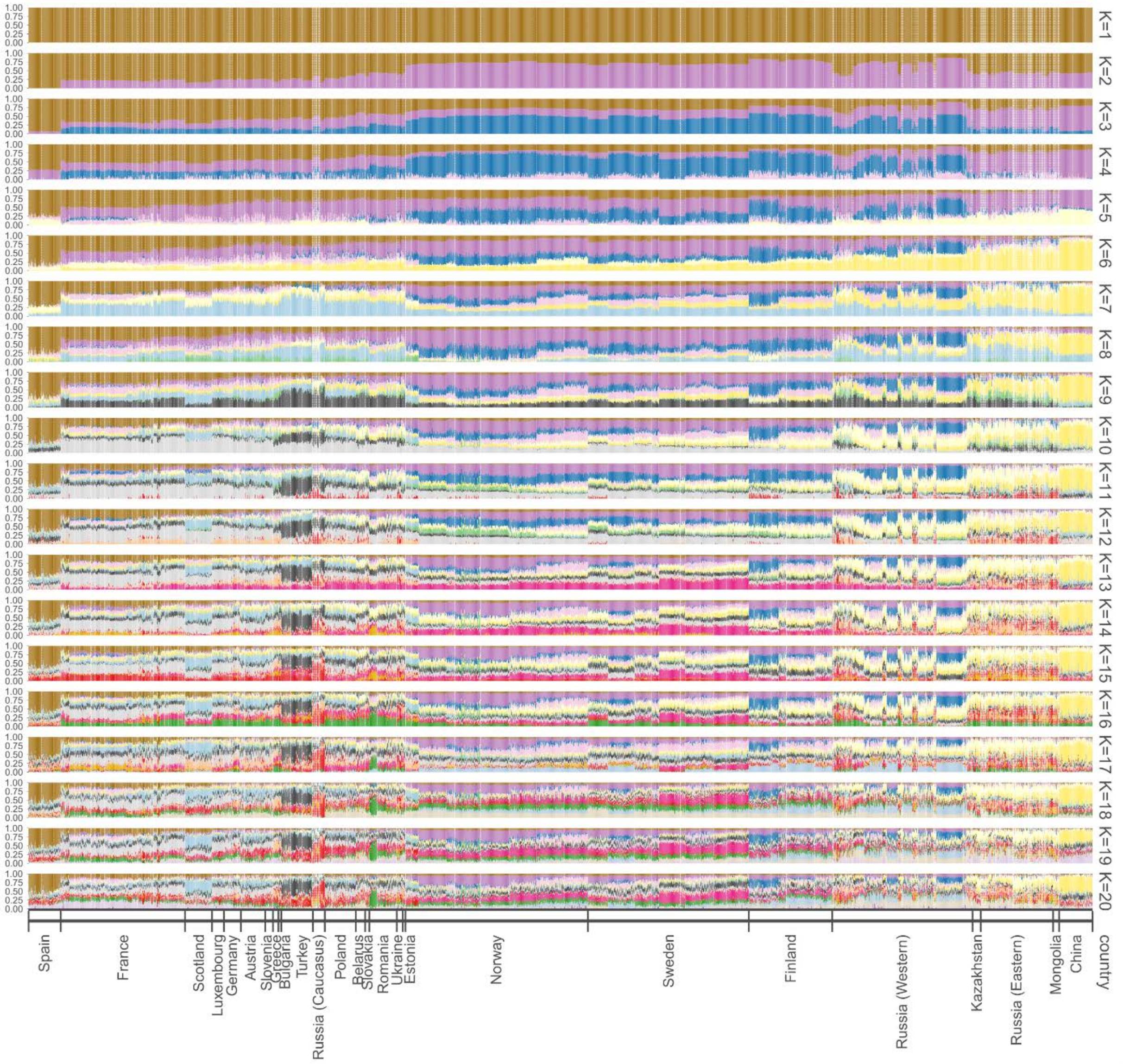


**Figure S9**: Admixture inference produced by tess3 and showing the proportion of each cluster component for *K* varying between 1 and 20 groups. The populations are ordered by country following the PCA results: from Spain to Scandinavia then to China. In each country, the populations are ordered by longitude except for Scandinavia where they are ordered by latitude.

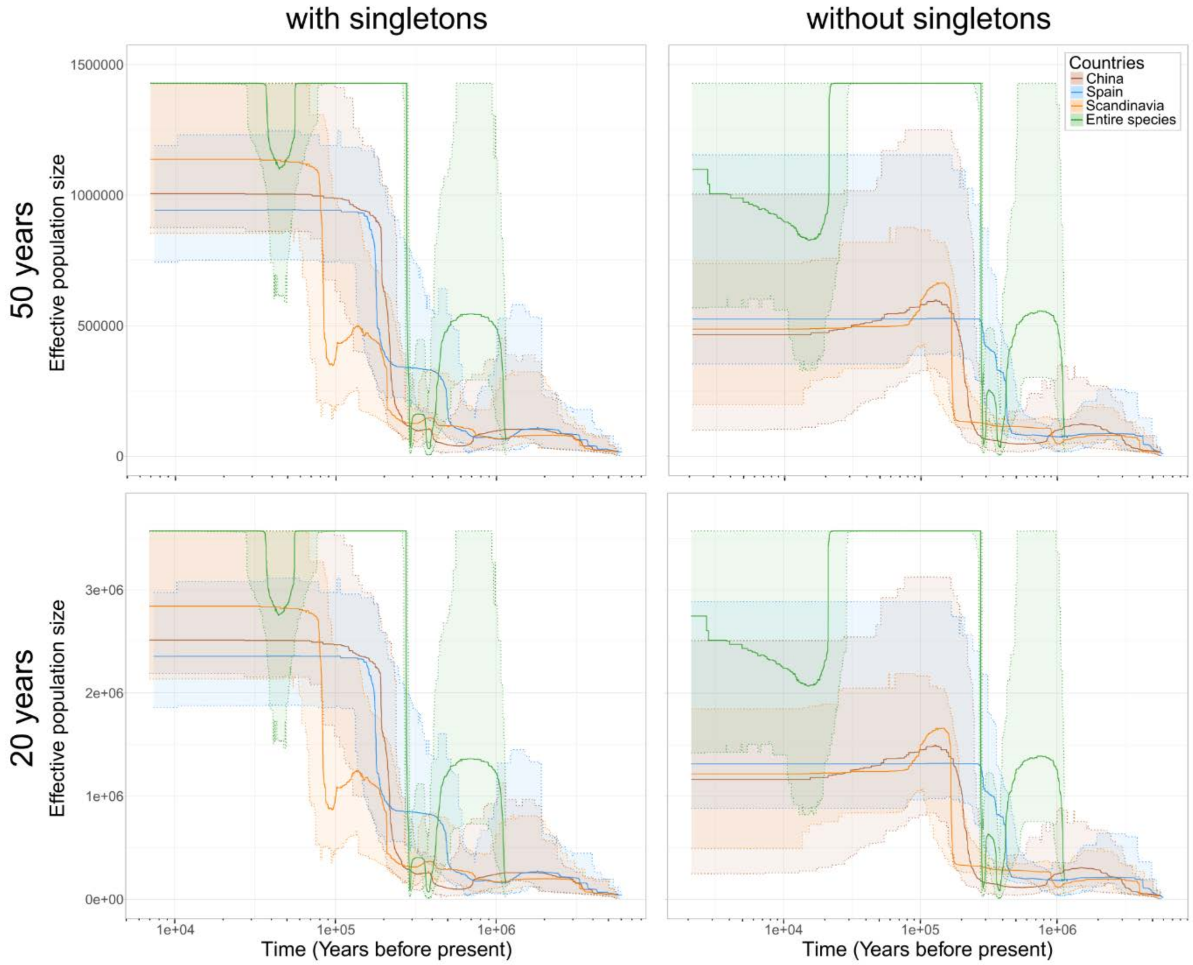


**Figure S10**: Population history of the three main *P. sylvestris* lineages and of the entire species. Generation times and theta variations are translated to years and effective population sizes by using a mutation rate of $7.10^{-10}$ mutations /site /year and a generation time of 50 years for the top row and 20 years for the bottom row. The plain lines represent the estimated median effective population sizes, and the shaded areas correspond to the 95% confidence interval. We used datasets with and without singletons, represented on the left and right respectively.

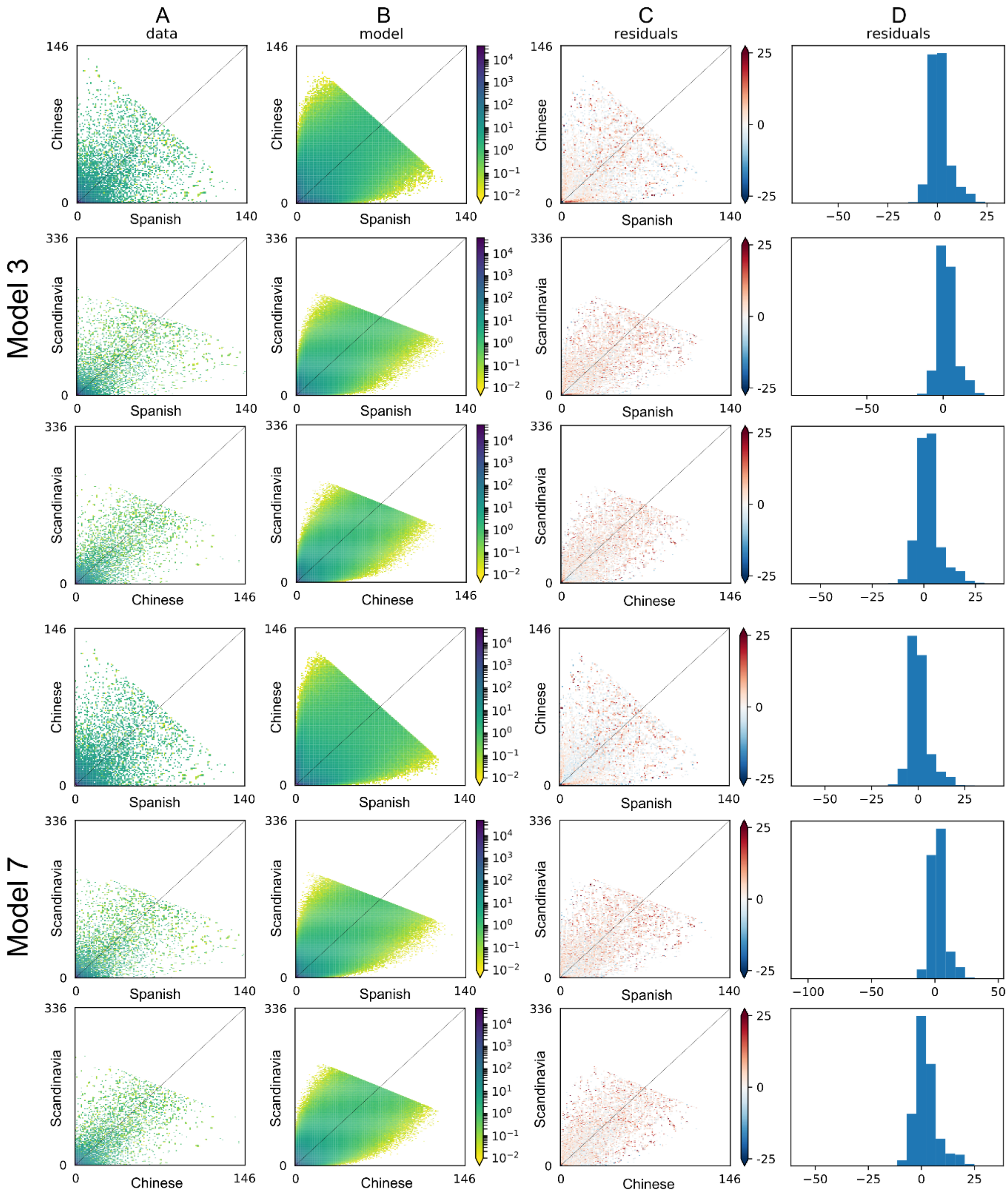


**Figure S11**: Goodness of fit of the model 3 and 7 inferred by fastsimcoal2. For each pair of groups, Spanish vs. Chinese (top), Spanish vs. Scandinavia (middle) and Chinese vs. Scandinavia (bottom), the observed joint folded SFS (data, A) is compared with the expected joint folded SFS (model, B), and the residuals between the data and the model are plotted in a colormap (C) and a histogram (D). In the colormap (C), red or blue residuals indicate that the model predicts too many or too few alleles in a given cell, respectively.

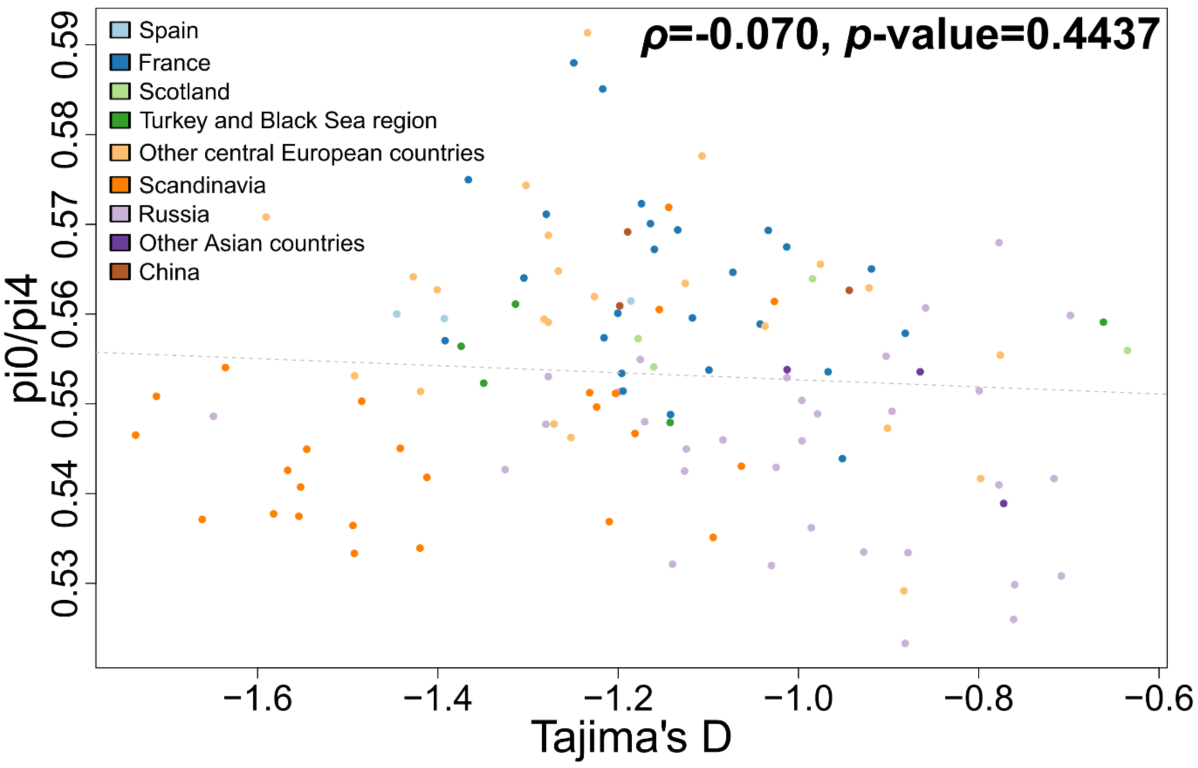


**Figure S12**: Variation of the $\pi_0/\pi_4$ ratio with Tajima's D values. Points represent large populations and are colored by regions. The regression lines (grey doted lines) are presented for illustration only, as the residuals of the linear model did not follow a normal distribution with homoscedasticity.

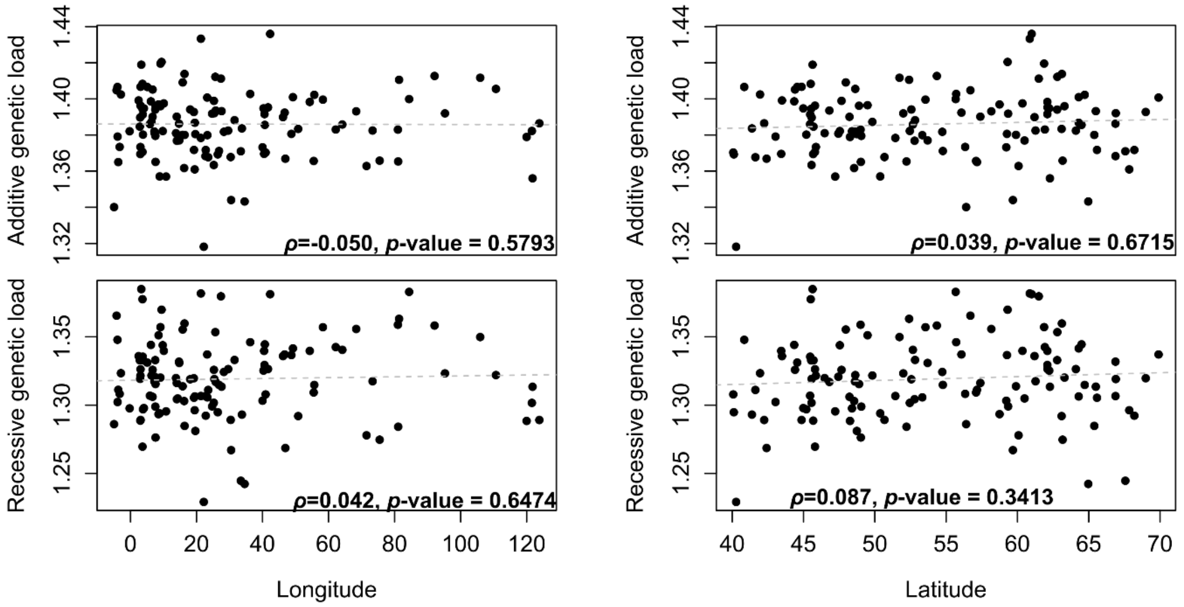


**Figure S13**: Additive (upper row) and recessive (lower row) genetic load in relation to longitude and latitude. Each point corresponds to one large population. The Spearman correlation coefficients and the p-values are indicated for each figure. The regression lines (grey doted lines) are presented for illustration only, as the residuals of the linear model did not follow a normal distribution with homoscedasticity.

#### Supplementary tables

Table S1: Pairwise nucleotide diversity, heterozygosity and inbreeding coefficient for each population, country or continental region. For each population, the coordinates (latitude and longitude) and the number of individuals ($N_{original}$) are given in the first columns. For populations smaller than 6 individuals, we merged several neighboring locations and give the total number of individuals analyzed ($N_{merged}$). The pairwise nucleotide diversity was calculated on the full dataset including invariant sites ($\pi$), and the SNPs were later restricted to the 0-fold ($\pi_0$) and 4-fold ($\pi_4$) sites. We calculated the inbreeding coefficient $F_{IS}$, the observed and expected heterozygosities ($H_o$ and $H_e$) on the dataset filtered for MAF 0.05. Six individuals from three very distant Russian populations were not included in these analyses. The values presented for countries and continental regions are averages of the individual populations results, weighted for the number of individuals per population.

| Original population name | Latitude | Longitude | $N_{original}$ | Merged population name | $N_{merged}$ | $\pi$ | $\pi_4$ | $\pi_0$ | $H_o$ | $H_e$ | $F_{IS}$ | Tajima's D |
|---|---|---|---|---|---|---|---|---|---|---|---|---|
| Whole distribution | | | | | 2315 | 0.00622 | 0.00929 | 0.00474 | 0.258 | 0.277 | 0.057 | -1.297 |
| Spain | | | | | 70 | 0.00631 | 0.00941 | 0.00489 | 0.265 | 0.273 | 0.027 | -1.369 |
| Embalse del Ebro | 43.03 | -3.91 | 15 | | | 0.00636 | 0.00949 | 0.00491 | 0.278 | 0.286 | 0.020 | -1.186 |
| Montana Soriano Burgalesa | 41.95 | -2.9 | 27 | | | 0.00631 | 0.00936 | 0.00490 | 0.262 | 0.270 | 0.029 | -1.446 |
| Sierra de Guadarrama | 40.84 | -3.94 | 28 | | | 0.00628 | 0.00943 | 0.00487 | 0.262 | 0.270 | 0.028 | -1.393 |
| Scotland | | | | | 59 | 0.00638 | 0.00942 | 0.00490 | 0.284 | 0.292 | 0.020 | -1.032 |
| Abernethy | 56.34 | -3.26 | 18 | | | 0.00642 | 0.00963 | 0.00496 | 0.276 | 0.283 | 0.018 | -1.161 |
| Glen Orchy | 56.4 | -4.97 | 11 | | | 0.00605 | 0.00882 | 0.00461 | 0.301 | 0.315 | 0.026 | -0.635 |
| Glenmore | 57.17 | -3.7 | 18 | | | 0.00643 | 0.00950 | 0.00493 | 0.273 | 0.283 | 0.028 | -1.178 |
| Rannoch | 56.7 | -4.2 | 12 | | | 0.00654 | 0.00953 | 0.00502 | 0.298 | 0.300 | 0.002 | -0.984 |
| Western and Central Europe | | | | | 526 | 0.00650 | 0.00967 | 0.00503 | 0.282 | 0.290 | 0.016 | -1.191 |
| France | | | | | 271 | 0.00655 | 0.00982 | 0.00512 | 0.284 | 0.293 | 0.018 | -1.158 |
| Allos | 44.43 | 6.71 | 8 | | | 0.00643 | 0.00950 | 0.00502 | 0.309 | 0.310 | -0.014 | -0.919 |
| Aydat | 45.68 | 2.99 | 10 | | | 0.00695 | 0.01008 | 0.00550 | 0.306 | 0.299 | -0.030 | -1.249 |
| Ayguebonne | 45.8 | 3.65 | 9 | | | 0.00663 | 0.01018 | 0.00536 | 0.279 | 0.304 | 0.052 | -1.134 |
| Bitche | 49.05 | 7.53 | 9 | | | 0.00686 | 0.01019 | 0.00544 | 0.312 | 0.300 | -0.044 | -1.174 |
| Bruyeres | 48.22 | 6.7 | 8 | | | 0.00681 | 0.01016 | 0.00537 | 0.324 | 0.308 | -0.055 | -1.013 |
| Chaudes-Aigues | 44.86 | 3.06 | 9 | | | 0.00652 | 0.00979 | 0.00520 | 0.319 | 0.300 | -0.055 | -0.967 |
| Cheire de Volvic | 45.87 | 3.02 | 9 | | | 0.00624 | 0.00931 | 0.00473 | 0.289 | 0.307 | 0.034 | -0.882 |
| Entremont le Vieux | 45.46 | 5.91 | 13 | | | 0.00640 | 0.00952 | 0.00496 | 0.274 | 0.285 | 0.022 | -1.142 |
| Kiffis | 47.44 | 7.36 | 9 | | | 0.00650 | 0.00966 | 0.00486 | 0.301 | 0.300 | -0.016 | -1.034 |
| Laussonne | 44.98 | 4.06 | 10 | | | 0.00659 | 0.00996 | 0.00527 | 0.302 | 0.294 | -0.029 | -1.099 |
| Malvieille | 45.5 | 3.64 | 13 | | | 0.00663 | 0.01035 | 0.00527 | 0.254 | 0.288 | 0.086 | -1.366 |
| Arlanc | 45.41 | 3.7 | 5 | Massif Central | 13 | 0.00667 | 0.00971 | 0.00515 | 0.286 | 0.295 | 0.021 | -1.195 |
| Massif Central | 44.88 | 3.7 | 8 | | | | | | | | | |

| Original population name | Latitude | Longitude | $N_{original}$ | Merged population name | $N_{merged}$ | $\pi$ | $\pi_4$ | $\pi_0$ | $H_o$ | $H_e$ | $F_{IS}$ | Tajima's D |
|---|---|---|---|---|---|---|---|---|---|---|---|---|
| Chanat | 45.83 | 2.98 | 5 | Massif Central 2 | 15 | 0.00629 | 0.00947 | 0.00486 | 0.237 | 0.279 | 0.114 | -1.305 |
| Fontfreyde | 45.71 | 2.98 | 10 | | | | | | | | | |
| Massif de la Comte | 45.63 | 3.28 | 10 | | | 0.00650 | 0.00999 | 0.00519 | 0.264 | 0.299 | 0.079 | -1.164 |
| Nohanent | 45.8 | 3.04 | 12 | | | 0.00641 | 0.00953 | 0.00493 | 0.254 | 0.288 | 0.087 | -1.200 |
| Haguenau | 48.85 | 7.71 | 5 | Northern Alsace | 12 | 0.00650 | 0.00966 | 0.00497 | 0.283 | 0.282 | -0.013 | -1.196 |
| Wimmenau | 48.93 | 7.44 | 7 | | | | | | | | | |
| Philippsbourg | 49.02 | 7.59 | 10 | | | 0.00659 | 0.00967 | 0.00511 | 0.302 | 0.296 | -0.025 | -1.118 |
| Le Fugeret | 44.03 | 6.67 | 5 | Southern Alps | 12 | 0.00663 | 0.00971 | 0.00508 | 0.309 | 0.306 | -0.012 | -1.073 |
| Luz-la-Croix-Haute | 44.67 | 5.71 | 7 | | | | | | | | | |
| Mont Sainte-Odile | 48.43 | 7.41 | 5 | Southern Alsace | 13 | 0.00668 | 0.00983 | 0.00513 | 0.286 | 0.282 | -0.016 | -1.280 |
| Walscheid | 48.6 | 7.23 | 8 | | | | | | | | | |
| St Amans Soult | 43.49 | 2.47 | 10 | | | 0.00677 | 0.01039 | 0.00540 | 0.300 | 0.295 | -0.023 | -1.160 |
| St Martin de Tours | 45.7 | 2.83 | 12 | | | 0.00634 | 0.00962 | 0.00496 | 0.256 | 0.286 | 0.075 | -1.216 |
| Tartaret | 45.57 | 2.93 | 10 | | | 0.00630 | 0.00953 | 0.00485 | 0.271 | 0.293 | 0.050 | -1.043 |
| Bollene | 44.32 | 4.77 | 3 | Vallée Rhône | 11 | 0.00685 | 0.00983 | 0.00539 | 0.296 | 0.291 | -0.012 | -1.217 |
| Die | 44.79 | 5.37 | 8 | | | | | | | | | |
| Villepail | 48.4 | -0.26 | 9 | | | 0.00635 | 0.00972 | 0.00494 | 0.300 | 0.295 | -0.023 | -0.951 |
| Virennes | 45.53 | 3.64 | 15 | | | 0.00653 | 0.01014 | 0.00522 | 0.254 | 0.278 | 0.064 | -1.392 |
| Luxembourg | | | | | | | | | | | | |
| Luxembourg | 49.82 | 6.35 | 26 | | | 0.00640 | 0.00948 | 0.00489 | 0.258 | 0.268 | 0.033 | -1.492 |
| Germany | | | | | 37 | 0.00663 | 0.00962 | 0.00502 | 0.278 | 0.287 | 0.019 | -1.306 |
| Bad Freienwalde | 52.79 | 14.05 | 8 | | | 0.00682 | 0.00988 | 0.00517 | 0.314 | 0.309 | -0.028 | -1.126 |
| Teufelsee, Echzell | 50.38 | 8.85 | 10 | | | 0.00687 | 0.00970 | 0.00526 | 0.299 | 0.302 | -0.004 | -1.234 |
| Darmstadt | 49.87 | 8.65 | 4 | Western Germany | 19 | 0.00642 | 0.00947 | 0.00484 | 0.252 | 0.270 | 0.051 | -1.419 |
| Karlsruhe | 49 | 8.4 | 12 | | | | | | | | | |
| Lampertheim | 49.6 | 8.47 | 3 | | | | | | | | | |
| Austria | | | | | 53 | 0.00643 | 0.00942 | 0.00491 | 0.285 | 0.292 | 0.017 | -1.125 |
| WKie 1 | 47.96 | 15.82 | 13 | | | 0.00646 | 0.00916 | 0.00490 | 0.285 | 0.294 | 0.018 | -1.107 |
| WKie 12 | 47.54 | 9.77 | 8 | | | 0.00647 | 0.00944 | 0.00493 | 0.326 | 0.314 | -0.043 | -0.922 |
| WKie 2 | 48.55 | 16.26 | 12 | | | 0.00652 | 0.00971 | 0.00503 | 0.303 | 0.302 | -0.009 | -1.037 |
| WKie 3 | 47.23 | 10.79 | 20 | | | 0.00635 | 0.00942 | 0.00484 | 0.257 | 0.277 | 0.057 | -1.271 |
| Slovenia | | | | | | | | | | | | |
| Kranjska Gora | 46.48 | 13.78 | 17 | | | 0.00641 | 0.00952 | 0.00486 | 0.278 | 0.280 | 0.007 | -1.252 |
| Slovakia | | | | | | | | | | | | |
| Cierny Balog | 48.67 | 19.67 | 9 | | | 0.00634 | 0.00941 | 0.00491 | 0.310 | 0.300 | -0.036 | -0.976 |
| Poland | | | | | 68 | 0.00641 | 0.00949 | 0.00496 | 0.266 | 0.277 | 0.026 | -1.325 |
| Dzisna | 53.7 | 14.78 | 16 | | | 0.00643 | 0.00958 | 0.00494 | 0.271 | 0.279 | 0.020 | -1.278 |
| Jedlnia | 51.44 | 21.32 | 14 | | | 0.00648 | 0.00944 | 0.00507 | 0.272 | 0.284 | 0.027 | -1.277 |
| Ostrowo | 52.56 | 18.08 | 17 | | | 0.00636 | 0.00950 | 0.00494 | 0.265 | 0.277 | 0.033 | -1.282 |
| Wilcza Jama | 53.33 | 23.29 | 21 | | | 0.00638 | 0.00943 | 0.00493 | 0.260 | 0.269 | 0.024 | -1.427 |
| Belarus | | | | | 20 | 0.00621 | 0.00960 | 0.00478 | 0.299 | 0.296 | -0.018 | -0.892 |

| Original population name | Latitude | Longitude | $N_{original}$ | Merged population name | $N_{merged}$ | $\pi$ | $\pi_4$ | $\pi_0$ | $H_o$ | $H_e$ | $F_{IS}$ | Tajima's D |
|---|---|---|---|---|---|---|---|---|---|---|---|---|
| Cheremshitsy | 54.77 | 26.83 | 10 | | | 0.00619 | 0.00938 | 0.00477 | 0.297 | 0.296 | -0.012 | -0.901 |
| Domzharytsy | 54.73 | 28.3 | 10 | | | 0.00624 | 0.00982 | 0.00480 | 0.302 | 0.296 | -0.024 | -0.883 |
| Estonia | | | | | | | | | | | | |
| Tallinn | 59.35 | 24.73 | 6 | | | 0.00633 | 0.00946 | 0.00484 | 0.346 | 0.325 | -0.072 | -0.776 |
| Bulgaria | | | | | | | | | | | | |
| Pirin | 41.62 | 23.55 | 6 | | | 0.00632 | 0.00954 | 0.00481 | 0.353 | 0.329 | -0.076 | -0.798 |
| Ukraine | | | | | | | | | | | | |
| Kiev P4 | 50.68 | 30.33 | 5 | Kiev | 13 | 0.00669 | 0.00986 | 0.00519 | 0.270 | 0.288 | 0.045 | -1.401 |
| Kiev P5 | 50.67 | 30.38 | 8 | | | | | | | | | |
| Southeastern Europe | | | | | 140 | 0.00644 | 0.00951 | 0.00496 | 0.267 | 0.275 | 0.024 | -1.356 |
| Greece | | | | | | | | | | | | |
| mount Pieria | 40.27 | 22.2 | 12 | | | 0.00709 | 0.01024 | 0.00548 | 0.324 | 0.304 | -0.056 | -1.302 |
| Romania | | | | | 60 | 0.00641 | 0.00957 | 0.00499 | 0.261 | 0.274 | 0.039 | -1.377 |
| Breaza | 47.68 | 25.22 | 23 | | | 0.00668 | 0.00969 | 0.00514 | 0.260 | 0.270 | 0.031 | -1.591 |
| Cheile Bicazului | 46.81 | 25.82 | 20 | | | 0.00624 | 0.00927 | 0.00481 | 0.260 | 0.272 | 0.034 | -1.226 |
| Piatra Craiului NP | 45.56 | 25.24 | 17 | | | 0.00623 | 0.00976 | 0.00499 | 0.262 | 0.282 | 0.057 | -1.266 |
| Turkey | | | | | 68 | 0.00635 | 0.00933 | 0.00485 | 0.263 | 0.271 | 0.026 | -1.347 |
| Canakkale | 40.11 | 26.44 | 21 | | | 0.00642 | 0.00931 | 0.00490 | 0.268 | 0.275 | 0.020 | -1.314 |
| Ilgaz | 40.92 | 33.69 | 18 | Central Turkey | 24 | 0.00625 | 0.00924 | 0.00478 | 0.255 | 0.268 | 0.041 | -1.349 |
| Kastamonu | 41.38 | 33.75 | 6 | | | | | | | | | |
| Eskipolat | 40.07 | 40.93 | 23 | | | 0.00638 | 0.00944 | 0.00489 | 0.266 | 0.271 | 0.015 | -1.374 |
| Eastern Europe and Asia | | | | | 589 | 0.00610 | 0.00917 | 0.00465 | 0.268 | 0.283 | 0.045 | -1.102 |
| Russia | | | | | 485 | 0.00613 | 0.00924 | 0.00466 | 0.266 | 0.285 | 0.051 | -1.109 |
| Krestyanka, Altai | 52.47 | 81.63 | 3 | Altai | 12 | 0.00608 | 0.00925 | 0.00455 | 0.284 | 0.292 | 0.014 | -1.140 |
| Mamontovo, Altai | 52.7 | 81.63 | 3 | | | | | | | | | |
| Mikhaylovsky District, Altai | 51.82 | 79.78 | 2 | | | | | | | | | |
| Novichikha, Altai | 52.21 | 81.38 | 2 | | | | | | | | | |
| Srosty, Altai | 51.9 | 81 | 2 | | | | | | | | | |
| Krugloye, Altai | 51.32 | 80.37 | 2 | Altai 2 | 14 | 0.00635 | 0.00955 | 0.00495 | 0.277 | 0.289 | 0.028 | -0.996 |
| Makarovo, Altai | 53.28 | 82 | 3 | | | | | | | | | |
| Podstepnoye, Altai | 52.97 | 82.1 | 3 | | | | | | | | | |
| Poselok Peresheyechnyy, Altai | 51.73 | 80.87 | 1 | | | | | | | | | |
| Rebrikha, Altai | 53.07 | 82.37 | 2 | | | | | | | | | |
| Volchikha, Altai | 52.05 | 80.43 | 3 | | | | | | | | | |
| Apatity, Murmansk | 67.57 | 33.43 | 6 | | | 0.00639 | 0.00954 | 0.00491 | 0.362 | 0.333 | -0.088 | -0.777 |
| Archangels plantation | 61 | 42.3 | 24 | | | 0.00621 | 0.00932 | 0.00470 | 0.239 | 0.271 | 0.098 | -1.278 |
| Archangels stand 13 | 64.5 | 40.7 | 26 | | | 0.00597 | 0.00887 | 0.00453 | 0.227 | 0.269 | 0.128 | -1.198 |
| Archangels stand 3 | 61.2 | 46.2 | 20 | | | 0.00605 | 0.00898 | 0.00451 | 0.243 | 0.270 | 0.076 | -1.171 |

| Original population name | Latitude | Longitude | $N_{original}$ | Merged population name | $N_{merged}$ | $\pi$ | $\pi_4$ | $\pi_0$ | $H_o$ | $H_e$ | $F_{IS}$ | Tajima's D |
|---|---|---|---|---|---|---|---|---|---|---|---|---|
| Archangels stand 5 | 62.1 | 40.6 | 14 | | | 0.00613 | 0.00910 | 0.00458 | 0.237 | 0.285 | 0.131 | -1.084 |
| Konosha, Arkhangelsk | 60.92 | 40.8 | 7 | Arkhangelsk | 16 | 0.00604 | 0.00921 | 0.00456 | 0.276 | 0.274 | -0.006 | -0.986 |
| Plesetsk, Arkhangelsk | 63.33 | 41.08 | 2 | | | | | | | | | |
| Shalakusha, Arkhangelsk | 62.22 | 40.27 | 7 | | | | | | | | | |
| Arshan, Irkutsk | 51.9 | 102.45 | 4 | Baikal | 16 | 0.00611 | 0.00927 | 0.00475 | 0.278 | 0.288 | 0.023 | -1.013 |
| Djidinksi district, Bouryatia | 50.83 | 105.95 | 1 | | | | | | | | | |
| Ivolginsky District, Buryatia | 51.8 | 107.27 | 7 | | | | | | | | | |
| Olkhon Island, Irkutsk | 53.13 | 107.38 | 2 | | | | | | | | | |
| Selenga river mouth, Buryatia | 50.98 | 106.6 | 2 | | | | | | | | | |
| Achishkho, Krasnodar | 43.72 | 40.18 | 3 | Black Sea | 17 | 0.00616 | 0.00924 | 0.00486 | 0.240 | 0.281 | 0.114 | -1.142 |
| Guzeripl, Adygea | 44 | 40.13 | 2 | | | | | | | | | |
| Kars, Turkey | 40.72 | 42.55 | 1 | | | | | | | | | |
| Psebay, Krasnodar | 43.98 | 40.8 | 1 | | | | | | | | | |
| Ritsa, Abkhazia | 43.47 | 40.53 | 5 | | | | | | | | | |
| Rosa Khutor, Krasnodar | 43.68 | 40.28 | 1 | | | | | | | | | |
| Teberda, Karachay-Cherkess | 43.38 | 41.72 | 2 | | | | | | | | | |
| Utrish island, Krasnodar | 44.77 | 37.42 | 2 | | | | | | | | | |
| Bolotnoye, Bolotninsky | 55.67 | 84.4 | 7 | | | 0.00612 | 0.00972 | 0.00481 | 0.332 | 0.322 | -0.042 | -0.760 |
| Aneyeva, Khanty-Mansiysk | 63.15 | 63.97 | 6 | Eastern Ural | 14 | 0.00619 | 0.00932 | 0.00467 | 0.278 | 0.288 | 0.021 | -1.126 |
| Ivdel, Sverdlovsk | 60.7 | 60.38 | 8 | | | | | | | | | |
| Gunibsky District, Daghestan | 42.41 | 46.91 | 10 | | | 0.00599 | 0.00879 | 0.00456 | 0.327 | 0.314 | -0.042 | -0.662 |
| Kem, Karelia | 64.97 | 34.57 | 8 | | | 0.00628 | 0.00945 | 0.00481 | 0.315 | 0.315 | -0.016 | -0.896 |
| Kadzherom, Komi | 64.72 | 55.68 | 2 | Komi | 10 | 0.00601 | 0.00924 | 0.00455 | 0.275 | 0.294 | 0.043 | -0.879 |
| Sosnogorsk, Komi | 63.8 | 54.8 | 8 | | | | | | | | | |
| Megdurechensk, Komi | 63.1 | 50.8 | 16 | | | 0.00599 | 0.00897 | 0.00446 | 0.248 | 0.277 | 0.081 | -0.979 |
| Navlinsky District, Bryansk | 52.82 | 31.52 | 6 | | | 0.00635 | 0.00975 | 0.00479 | 0.328 | 0.337 | 0.000 | -0.761 |
| Noyabrsk, Yamalo-Nenets | 63.17 | 75.47 | 9 | | | 0.00599 | 0.00893 | 0.00459 | 0.299 | 0.299 | -0.012 | -0.799 |
| Obozersky, Arkhangelsk | 62.23 | 40.3 | 8 | | | 0.00611 | 0.00925 | 0.00466 | 0.310 | 0.299 | -0.041 | -0.716 |
| Pomozdino, Komi | 62.1 | 54.3 | 59 | | | 0.00606 | 0.00902 | 0.00456 | 0.225 | 0.261 | 0.123 | -1.649 |
| Mirny, Sakha | 62.53 | 113.98 | 1 | Sakha | 7 | 0.00597 | 0.00876 | 0.00448 | 0.311 | 0.330 | 0.031 | -0.698 |
| Yakutsk, Sakha | 62.03 | 129.6 | 6 | | | | | | | | | |

| Original population name | Latitude | Longitude | $N_{original}$ | Merged population name | $N_{merged}$ | $\pi$ | $\pi_4$ | $\pi_0$ | $H_o$ | $H_e$ | $F_{IS}$ | Tajima's D |
|---|---|---|---|---|---|---|---|---|---|---|---|---|
| Salym, Khanty-Mansiysk | 60.08 | 71.5 | 9 | | | 0.00622 | 0.00930 | 0.00479 | 0.302 | 0.302 | -0.012 | -0.903 |
| Alandskoye, Orenburg | 52.17 | 59.73 | 2 | South Ural | 23 | 0.00624 | 0.00953 | 0.00482 | 0.254 | 0.270 | 0.053 | -1.325 |
| Ay, Bashkortostan | 55.97 | 58.2 | 1 | | | | | | | | | |
| Birsk, Bashkortostan | 55.33 | 55.6 | 2 | | | | | | | | | |
| Buzuluk, Orenburg | 52.77 | 52.27 | 2 | | | | | | | | | |
| Chebarkulsky District, Chelyabinsk | 54.6 | 60.7 | 2 | | | | | | | | | |
| Chesmensky District, Chelyabinsk | 53.88 | 60.45 | 2 | | | | | | | | | |
| Inzer, Bashkortostan | 54.23 | 57.5 | 1 | | | | | | | | | |
| Isjkildino, Bashkortostan | 53.43 | 58.27 | 2 | | | | | | | | | |
| Karagay-Pokrovka, Orenburg | 51.63 | 57.9 | 2 | | | | | | | | | |
| Kropachevo, Chelyabinsk | 54.98 | 57.95 | 1 | | | | | | | | | |
| Kvarkensky District, Orenburg | 52.22 | 60 | 2 | | | | | | | | | |
| Parij, Chelyabinsk | 53.27 | 60.03 | 2 | | | | | | | | | |
| Peschanka, Chelyabinsk | 52.98 | 59.87 | 1 | | | | | | | | | |
| Zilair, Bashkortostan | 52.25 | 57.53 | 1 | | | | | | | | | |
| Pavlovsk, St. Petersburg | 59.68 | 30.48 | 1 | St. Petersburg | 17 | 0.00637 | 0.00968 | 0.00489 | 0.268 | 0.278 | 0.025 | -1.280 |
| Pskov, Pskov | 57.82 | 28.55 | 2 | | | | | | | | | |
| South Karelia | 61.78 | 33.81 | 4 | | | | | | | | | |
| Tikhvin, Leningrad | 59.67 | 33.5 | 10 | | | | | | | | | |
| Surgut, Khanty-Mansiysk | 61.27 | 73.33 | 9 | | | 0.00615 | 0.00894 | 0.00469 | 0.306 | 0.304 | -0.016 | -0.859 |
| Minskiy Pereulok, Tuva | 52.12 | 95.98 | 5 | Tian Shan | 14 | 0.00600 | 0.00901 | 0.00460 | 0.275 | 0.289 | 0.035 | -0.882 |
| Shagonar, Tuva | 51.52 | 93.23 | 7 | | | | | | | | | |
| Todzha Lake, Tuva | 52.35 | 96.55 | 2 | | | | | | | | | |
| Aradan, Krasnoyarsk | 52.57 | 93.45 | 2 | Tian Shan 2 | 12 | 0.00622 | 0.00975 | 0.00476 | 0.264 | 0.299 | 0.088 | -0.996 |
| Bograd, Khakassia | 54.23 | 90.82 | 1 | | | | | | | | | |
| Minusinsk, Krasnoyarsk | 53.68 | 91.73 | 2 | | | | | | | | | |
| Pogorelka, Krasnoyarsk | 56.37 | 92.95 | 2 | | | | | | | | | |
| Shira, Khakassia | 54.4 | 89.98 | 2 | | | | | | | | | |
| Tanzybey, Krasnoyarsk | 53.07 | 92.57 | 2 | | | | | | | | | |
| Yesaulovo, Krasnoyarsk | 56.13 | 93.25 | 1 | | | | | | | | | |

| Original population name | Latitude | Longitude | $N_{original}$ | Merged population name | $N_{merged}$ | $\pi$ | $\pi_4$ | $\pi_0$ | $H_o$ | $H_e$ | $F_{IS}$ | Tajima's D |
|---|---|---|---|---|---|---|---|---|---|---|---|---|
| Tobolsk, Tyumen | 58.17 | 68.37 | 10 | | | 0.00607 | 0.00912 | 0.00455 | 0.293 | 0.299 | 0.004 | -0.928 |
| Tyumen oblast | 57.1 | 55.5 | 8 | | | 0.00606 | 0.00928 | 0.00464 | 0.310 | 0.304 | -0.031 | -0.778 |
| Udora, Komi | 64.3 | 49.2 | 17 | | | 0.00596 | 0.00897 | 0.00452 | 0.239 | 0.278 | 0.113 | -1.025 |
| Bor, Nizhny Novgorod | 56.37 | 44.15 | 7 | Volga | 15 | 0.00630 | 0.00951 | 0.00487 | 0.274 | 0.281 | 0.016 | -1.175 |
| Kirya, Chuvash | 55.15 | 46.92 | 4 | | | | | | | | | |
| Kokshaysk, Mari El | 56.15 | 47.82 | 2 | | | | | | | | | |
| Yoshkar-Ola, Mari El | 56.62 | 47.92 | 2 | | | | | | | | | |
| Vologda, Vologda | 59.22 | 40 | 8 | | | 0.00605 | 0.00923 | 0.00450 | 0.306 | 0.299 | -0.032 | -0.708 |
| Kalouga, Kalouga | 54.52 | 36.2 | 2 | Western Russia | 10 | 0.00637 | 0.00977 | 0.00489 | 0.284 | 0.293 | 0.014 | -1.030 |
| Kovrov, Vladimir | 56.33 | 41.35 | 1 | | | | | | | | | |
| Lesnoy, Smolensk | 55.22 | 31.5 | 6 | | | | | | | | | |
| Tver, Tver | 56.75 | 36 | 1 | | | | | | | | | |
| Kirs, Kirov | 59.27 | 52.45 | 2 | Western Russia 2 | 14 | 0.00631 | 0.00976 | 0.00491 | 0.273 | 0.286 | 0.034 | -1.124 |
| Kotelnich, Kirov | 58 | 48.88 | 2 | | | | | | | | | |
| Krasnoborsk, Arkhangelsk | 61.85 | 45.88 | 2 | | | | | | | | | |
| Kumyonsky District, Kirov | 58.17 | 49.93 | 1 | | | | | | | | | |
| Lalsk, Kirov | 60.63 | 47.27 | 3 | | | | | | | | | |
| Pyshchug, Kostroma | 58.85 | 45.63 | 2 | | | | | | | | | |
| Zuyevka, Kirov | 58.1 | 51.13 | 2 | | | | | | | | | |
| Baykit, Krasnoyarsk | 61.68 | 96.37 | 3 | | | | | | | | | |
| Boguchany, Krasnoyarsk | 58.37 | 97.45 | 2 | | | | | | | | | |
| Touroukhansk, Krasnoyarsk | 65.8 | 87.96 | 1 | | | | | | | | | |
| Kazakhstan | | | | | 18 | 0.00612 | 0.00930 | 0.00474 | 0.317 | 0.304 | -0.045 | -0.819 |
| Amankaragaj | 52.67 | 64.17 | 9 | | | 0.00622 | 0.00932 | 0.00480 | 0.316 | 0.303 | -0.046 | -0.865 |
| Semey | 49 | 81 | 9 | | | 0.00602 | 0.00928 | 0.00467 | 0.318 | 0.306 | -0.044 | -0.772 |
| Mongolia | | | | | | | | | | | | |
| Bayan-Adarga | 48.55 | 111.07 | 2 | Mongolia | 13 | 0.00616 | 0.00930 | 0.00484 | 0.264 | 0.297 | 0.085 | -1.013 |
| Bogd Khan Uul | 47.82 | 106.87 | 2 | | | | | | | | | |
| Ereenii Nuruu | 48.77 | 111.6 | 2 | | | | | | | | | |
| Ku | 48.75 | 111.6 | 3 | | | | | | | | | |
| Onon | 48.65 | 110.43 | 2 | | | | | | | | | |
| Ulikhan | 49.23 | 112.62 | 2 | | | | | | | | | |
| China | | | | | 73 | 0.00591 | 0.00868 | 0.00452 | 0.265 | 0.269 | 0.016 | -1.139 |
| Honghua'erjizhen | 48.26 | 119.99 | 30 | | | 0.00587 | 0.00868 | 0.00450 | 0.258 | 0.265 | 0.024 | -1.198 |
| Mehe | 52.45 | 121.58 | 27 | | | 0.00590 | 0.00856 | 0.00449 | 0.260 | 0.267 | 0.025 | -1.190 |
| Tieling | 42.27 | 123.87 | 16 | | | 0.00599 | 0.00888 | 0.00459 | 0.285 | 0.280 | -0.015 | -0.944 |
| Northern Europe | | | | | 931 | 0.00610 | 0.00911 | 0.00457 | 0.235 | 0.265 | 0.099 | -1.483 |

| Original population name | Latitude | Longitude | $N_{original}$ | Merged population name | $N_{merged}$ | $\pi$ | $\pi_4$ | $\pi_0$ | $H_o$ | $H_e$ | $F_{IS}$ | Tajima's D |
|---|---|---|---|---|---|---|---|---|---|---|---|---|
| Norway | | | | | 398 | 0.00607 | 0.00903 | 0.00450 | 0.236 | 0.265 | 0.100 | -1.488 |
| Alta, Stengelsen | 69.9 | 23.3 | 52 | | | 0.00594 | 0.00879 | 0.00432 | 0.200 | 0.259 | 0.200 | -1.493 |
| Beiarn, Nordland | 66.9 | 14.3 | 59 | | | 0.00597 | 0.00889 | 0.00441 | 0.230 | 0.260 | 0.106 | -1.552 |
| Evje og Hornnes - Gjerstad | 58.7 | 8.5 | 13 | | | 0.00657 | 0.00953 | 0.00504 | 0.297 | 0.293 | -0.015 | -1.144 |
| Hemne, Sor-Trondelag | 63.3 | 9 | 63 | | | 0.00598 | 0.00896 | 0.00442 | 0.236 | 0.260 | 0.089 | -1.554 |
| Kirkesmoen, Troms | 69 | 19.2 | 60 | | | 0.00597 | 0.00888 | 0.00442 | 0.210 | 0.258 | 0.163 | -1.582 |
| Molde, Gjemnes, Skodje | 62.8 | 7.5 | 56 | | | 0.00629 | 0.00936 | 0.00465 | 0.268 | 0.271 | 0.023 | -1.420 |
| Ringerike, Buskerud | 60.4 | 10 | 60 | | | 0.00604 | 0.00902 | 0.00449 | 0.245 | 0.261 | 0.061 | -1.567 |
| Vågå, Oppland | 61.5 | 9 | 19 | | | 0.00605 | 0.00886 | 0.00448 | 0.216 | 0.275 | 0.172 | -1.232 |
| Valesæter - Telemark | 59.1 | 9.6 | 16 | | | 0.00659 | 0.00980 | 0.00514 | 0.301 | 0.296 | -0.020 | -1.154 |
| Sweden | | | | | 351 | 0.00612 | 0.00921 | 0.00463 | 0.236 | 0.264 | 0.093 | -1.505 |
| Arvidsjaur | 65.58 | 19.2 | 33 | | | 0.00613 | 0.00915 | 0.00469 | 0.216 | 0.261 | 0.146 | -1.636 |
| Dala Jarna | 60.5 | 14.3 | 37 | | | 0.00602 | 0.00920 | 0.00454 | 0.230 | 0.261 | 0.102 | -1.494 |
| Gunnilbo, Kulheden | 59.9 | 15.8 | 58 | | | 0.00613 | 0.00905 | 0.00453 | 0.223 | 0.261 | 0.130 | -1.735 |
| Hammarstrand | 63.1 | 16.2 | 16 | | | 0.00616 | 0.00909 | 0.00462 | 0.233 | 0.280 | 0.132 | -1.181 |
| Isopuolamajarvi | 66.88 | 22.7 | 29 | | | 0.00605 | 0.00898 | 0.00458 | 0.217 | 0.261 | 0.140 | -1.485 |
| Kiruna | 67.84 | 19.46 | 29 | | | 0.00595 | 0.00913 | 0.00451 | 0.228 | 0.263 | 0.115 | -1.210 |
| Kuttainen | 68.2 | 22.8 | 48 | | | 0.00615 | 0.00932 | 0.00467 | 0.245 | 0.265 | 0.069 | -1.442 |
| Lillberget | 64.3 | 19.5 | 47 | | | 0.00608 | 0.00942 | 0.00469 | 0.257 | 0.265 | 0.034 | -1.412 |
| Skillingaryd | 57.4 | 14 | 44 | | | 0.00629 | 0.00932 | 0.00476 | 0.247 | 0.263 | 0.054 | -1.712 |
| Slussfors | 65.39 | 16.4 | 10 | | | 0.00646 | 0.00956 | 0.00495 | 0.303 | 0.291 | -0.038 | -1.027 |
| Finland | | | | | 182 | 0.00610 | 0.00909 | 0.00458 | 0.233 | 0.267 | 0.109 | -1.428 |
| Äänekoski | 62.8 | 25.7 | 17 | | | 0.00608 | 0.00917 | 0.00460 | 0.246 | 0.276 | 0.088 | -1.063 |
| Kuhmo | 64.1 | 29.6 | 62 | | | 0.00602 | 0.00902 | 0.00450 | 0.226 | 0.261 | 0.120 | -1.661 |
| Pudasjärvi | 65.5 | 27.6 | 19 | | | 0.00610 | 0.00901 | 0.00460 | 0.245 | 0.269 | 0.068 | -1.203 |
| Ristiina | 61.5 | 27.4 | 17 | | | 0.00618 | 0.00936 | 0.00464 | 0.253 | 0.281 | 0.075 | -1.095 |
| Rovaniemen mlk | 66.9 | 25.4 | 19 | | | 0.00612 | 0.00903 | 0.00459 | 0.237 | 0.271 | 0.101 | -1.224 |
| Uusikaupunki | 60.9 | 21.3 | 48 | | | 0.00616 | 0.00912 | 0.00463 | 0.225 | 0.264 | 0.135 | -1.545 |

Table S2: Description and source of the 74 variables examined for genotype environment association.

| Variable | Type | Description | Source | Ref. |
|---|---|---|---|---|
| ai | Climate | Aridity index | https://chelsa-climate.org/ | (Karger *et al.*, 2017) |
| bio1 | Climate | Mean annual air temperature | https://chelsa-climate.org/ | (Karger *et al.*, 2017) |
| bio2 | Climate | Mean diurnal air temperature range | https://chelsa-climate.org/ | (Karger *et al.*, 2017) |
| bio3 | Climate | Ratio of diurnal variation to annual variation in temperatures | https://chelsa-climate.org/ | (Karger *et al.*, 2017) |
| bio4 | Climate | Standard deviation of the monthly mean temperatures | https://chelsa-climate.org/ | (Karger *et al.*, 2017) |
| bio5 | Climate | Mean daily maximum air temperature of the warmest month | https://chelsa-climate.org/ | (Karger *et al.*, 2017) |
| bio6 | Climate | Mean daily minimum air temperature of the coldest month | https://chelsa-climate.org/ | (Karger *et al.*, 2017) |
| bio7 | Climate | Annual range of air temperature | https://chelsa-climate.org/ | (Karger *et al.*, 2017) |
| bio8 | Climate | Mean daily mean air temperatures of the wettest quarter | https://chelsa-climate.org/ | (Karger *et al.*, 2017) |
| bio9 | Climate | Mean daily mean air temperatures of the driest quarter | https://chelsa-climate.org/ | (Karger *et al.*, 2017) |
| bio10 | Climate | Mean daily mean air temperatures of the warmest quarter | https://chelsa-climate.org/ | (Karger *et al.*, 2017) |
| bio11 | Climate | Mean daily mean air temperatures of the coldest quarter | https://chelsa-climate.org/ | (Karger *et al.*, 2017) |
| bio12 | Climate | Annual precipitation amount | https://chelsa-climate.org/ | (Karger *et al.*, 2017) |
| bio13 | Climate | Precipitation amount of the wettest month | https://chelsa-climate.org/ | (Karger *et al.*, 2017) |
| bio14 | Climate | Precipitation amount of the driest month | https://chelsa-climate.org/ | (Karger *et al.*, 2017) |
| bio15 | Climate | Precipitation Seasonality (Coefficient of Variation) | https://chelsa-climate.org/ | (Karger *et al.*, 2017) |
| bio16 | Climate | Mean monthly precipitation amount of the wettest quarter | https://chelsa-climate.org/ | (Karger *et al.*, 2017) |
| bio17 | Climate | Mean monthly precipitation amount of the driest quarter | https://chelsa-climate.org/ | (Karger *et al.*, 2017) |
| bio18 | Climate | Mean monthly precipitation amount of the warmest quarter | https://chelsa-climate.org/ | (Karger *et al.*, 2017) |
| bio19 | Climate | Mean monthly precipitation amount of the coldest quarter | https://chelsa-climate.org/ | (Karger *et al.*, 2017) |
| gdd0 | Climate | heat sum of all days above the 0°C | https://chelsa-climate.org/ | (Karger *et al.*, 2017) |

| Variable | Type | Description | Source | Ref. |
|---|---|---|---|---|
| | | temperature accumulated over 1 year. | | |
| gdd5 | Climate | heat sum of all days above the 5°C temperature accumulated over 1 year. | https://chelsa-climate.org/ | (Karger *et al.*, 2017) |
| gdd10 | Climate | heat sum of all days above the 10°C temperature accumulated over 1 year. | https://chelsa-climate.org/ | (Karger *et al.*, 2017) |
| gddlgd10 | Climate | Last day of the year above 10°C | https://chelsa-climate.org/ | (Karger *et al.*, 2017) |
| gdgfgd10 | Climate | First day of the year above 10°C | https://chelsa-climate.org/ | (Karger *et al.*, 2017) |
| gsl | Climate | growing season length TREELIM | https://chelsa-climate.org/ | (Paulsen & Körner, 2014; Karger *et al.*, 2017) |
| gsp | Climate | precipitation sum accumulated on all days during the growing season based on TREELIM | https://chelsa-climate.org/ | (Karger *et al.*, 2017) |
| gst | Climate | Mean temperature of the growing season TREELIM | https://chelsa-climate.org/ | (Karger *et al.*, 2017) |
| kg0 | Climate | Köppen Geiger | https://chelsa-climate.org/ | (Köppen & Geiger, 1936; Karger *et al.*, 2017) |
| kg1 | Climate | Köppen Geiger without As/Aw differentiation | https://chelsa-climate.org/ | (Karger *et al.*, 2017) |
| kg2 | Climate | Köppen Geiger after Peel et al. 2007 | https://chelsa-climate.org/ | (Peel *et al.*, 2007; Karger *et al.*, 2017) |
| kg3 | Climate | Wissmann 1939 | https://chelsa-climate.org/ | (von Wissmann, 1939; Karger *et al.*, 2017) |
| kg4 | Climate | Thornthwaite 1931 | https://chelsa-climate.org/ | (Thornthwaite, 1931; Karger *et al.*, 2017) |
| kg5 | Climate | Troll-Pfaffen | https://chelsa-climate.org/ | (Troll & Paffen, 1964; Karger *et al.*, 2017) |
| ngd0 | Climate | Number of days at which tas > 0°C. tas: daily mean air temperatures at 2 metres from hourly ERA5 data | https://chelsa-climate.org/ | (Karger *et al.*, 2017) |
| ngd5 | Climate | Number of days at which tas > 5°C | https://chelsa-climate.org/ | (Karger *et al.*, 2017) |
| ngd10 | Climate | Number of days at which tas > 10°C | https://chelsa-climate.org/ | (Karger *et al.*, 2017) |
| npp | Climate | Net primary productivity | https://chelsa-climate.org/ | (Lieth, 1972; Karger *et al.*, 2017) |
| pet_1 | Climate | Potential evaporation of the 1st month | https://chelsa-climate.org/ | (Karger *et al.*, 2017) |
| pet_2 | Climate | Potential evaporation | https://chelsa-climate.org/ | (Karger *et al.*, 2017) |
| pet_3 | Climate | Potential evaporation | https://chelsa-climate.org/ | (Karger *et al.*, 2017) |
| pet_4 | Climate | Potential evaporation | https://chelsa-climate.org/ | (Karger *et al.*, 2017) |
| pet_5 | Climate | Potential evaporation | https://chelsa-climate.org/ | (Karger *et al.*, 2017) |
| pet_6 | Climate | Potential evaporation | https://chelsa-climate.org/ | (Karger *et al.*, 2017) |
| pet_7 | Climate | Potential evaporation | https://chelsa-climate.org/ | (Karger *et al.*, 2017) |

| Variable | Type | Description | Source | Ref. |
|---|---|---|---|---|
| pet_8 | Climate | Potential evaporation | https://chelsa-climate.org/ | (Karger *et al.*, 2017) |
| pet_9 | Climate | Potential evaporation | https://chelsa-climate.org/ | (Karger *et al.*, 2017) |
| pet_10 | Climate | Potential evaporation | https://chelsa-climate.org/ | (Karger *et al.*, 2017) |
| pet_11 | Climate | Potential evaporation | https://chelsa-climate.org/ | (Karger *et al.*, 2017) |
| scd | Climate | Number of days with snowcover calculated using the snowpack model implementation in from TREELIM | https://chelsa-climate.org/ | (Karger *et al.*, 2017) |
| consensus1 | Species composition | Evergreen/deciduous needleleaf trees | http://www.earthenv.org/landcover | (Tuanmu & Jetz, 2014) |
| consensus4 | Species composition | Mixed/other trees | http://www.earthenv.org/landcover | (Tuanmu & Jetz, 2014) |
| gdd | Climate | Growing degree days | https://sage.nelson.wisc.edu/data-and-models/atlas-of-the-biosphere/mapping-the-biosphere/ecosystems/ | (New *et al.*, 1999; New *et al.*, 2000) |
| elev | Climate | Elevation values | https://www.worldclim.org/ | (Fick & Hijmans, 2017) |
| frs | Climate | Ground-frost frequency | https://www.ipcc-data.org/ | (Stockhause *et al.*, 2019) |
| uvb1 | Light conditions | Annual mean UV-B | https://www.ufz.de/gluv/ | (Beckmann *et al.*, 2014) |
| uvb2 | Light conditions | UV-B seasonality | https://www.ufz.de/gluv/ | (Beckmann *et al.*, 2014) |
| uvb3 | Light conditions | Mean UV-B of highest month | https://www.ufz.de/gluv/ | (Beckmann *et al.*, 2014) |
| uvb4 | Light conditions | mean UV-B of lowest month | https://www.ufz.de/gluv/ | (Beckmann *et al.*, 2014) |
| uvb5 | Light conditions | Sum of monthly mean UV-B during highest quarter | https://www.ufz.de/gluv/ | (Beckmann *et al.*, 2014) |
| uvb6 | Light conditions | Sum of monthly mean UV-B during lowest quarter | https://www.ufz.de/gluv/ | (Beckmann *et al.*, 2014) |
| vap | Climate | Vapor pressure | https://www.ipcc-data.org/ | (Stockhause *et al.*, 2019) |
| wet | Climate | Wet-day frequency | https://www.ipcc-data.org/ | (Stockhause *et al.*, 2019) |
| bdod | Soil | Bulk density of the fine earth fraction (kg/dm³) | https://files.isric.org/soilgrids | (Poggio *et al.*, 2021) |
| cec | Soil | Cation exchange capacity of the soil [mmol(c)/kg] | https://files.isric.org/soilgrids | (Poggio *et al.*, 2021) |
| cfvo | Soil | Volumetric fraction of coarse fragments (cm3/dm3) | https://files.isric.org/soilgrids | (Poggio *et al.*, 2021) |
| clay | Soil | Proportion of clay particles in the fine earth fraction (0–2 μm) (g/kg) | https://files.isric.org/soilgrids | (Poggio *et al.*, 2021) |
| nitrogen | Soil | Total nitrogen (N) (g/kg) | https://files.isric.org/soilgrids | (Poggio *et al.*, 2021) |
| ocd | Soil | Organic carbon density (kg/m³) | https://files.isric.org/soilgrids | (Poggio *et al.*, 2021) |
| ocs | Soil | Organic carbon stocks (kg/m²) | https://files.isric.org/soilgrids | (Poggio *et al.*, 2021) |
| phh2o | Soil | Soil pH (pH x 10) | https://files.isric.org/soilgrids | (Poggio *et al.*, 2021) |
| sand | Soil | Proportion of sand particlesin the fine earth | https://files.isric.org/soilgrids | (Poggio *et al.*, 2021) |

| Variable | Type | Description | Source | Ref. |
|---|---|---|---|---|
| | | fraction (50–2000 μm) (g/kg) | | |
| silt | Soil | Proportion of silt particles in the fine earth fraction (2–50 μm) (g/kg) | https://files.isric.org/soilgrids | (Poggio *et al.*, 2021) |
| soc | Soil | Soil organic carbon content in the fine earth fraction (dg/kg) | https://files.isric.org/soilgrids | (Poggio *et al.*, 2021) |

Table S3: Relative likelihood and AIC of the different models shown in Figure S2 ordered by decreasing AIC. $N_{par}$: number of parameters in the model. AIC: Akaike criterion, calculated as $2*N_{par}-2*\ln(\text{likelihood})$. $\Delta_{AIC}$: difference between the AIC of the model and the best AIC. $\Delta_L$: difference between the estimated log-likelihood and the one obtained if there was a perfect fit between the expected and observed SFS (expected log-likelihood).

| Model | $N_{par}$ | AIC | $\Delta_{AIC}$ | Log-Likelihood | $\Delta_L$ |
|---|---|---|---|---|---|
| 7 | 25 | 5,158,033.56 | 0 | -1,120,041.90 | 43,332.49 |
| 3 | 25 | 5,159,825.92 | 1,792.36 | -1,120,431.11 | 43,721.70 |
| 11 | 25 | 5,160,099.64 | 2,066.08 | -1,120,490.54 | 43,781.14 |
| 14 | 21 | 5,163,677.79 | 5,644.23 | -1,121,269.27 | 44,559.86 |
| 18 | 17 | 5,166,534.28 | 8,500.72 | -1,121,891.28 | 45,181.88 |
| 17 | 15 | 5,171,495.20 | 13,461.64 | -1,122,969.40 | 46,259.99 |
| 19 | 17 | 5,172,304.47 | 14,270.90 | -1,123,144.26 | 46,434.86 |
| 4 | 28 | 5,176,331.97 | 18,298.41 | -1,124,014.05 | 47,304.64 |
| 10 | 18 | 5,177,318.52 | 19,284.96 | -1,124,232.61 | 47,523.21 |
| 8 | 28 | 5,177,603.98 | 19,570.42 | -1,124,290.26 | 47,580.85 |
| 12 | 28 | 5,181,737.16 | 23,703.60 | -1,125,187.77 | 48,478.36 |
| 1 | 15 | 5,182,662.27 | 24,628.71 | -1,125,394.30 | 48,684.90 |
| 2 | 18 | 5,182,886.05 | 24,852.49 | -1,125,441.59 | 48,732.18 |
| 16 | 13 | 5,183,738.00 | 25,704.44 | -1,125,628.76 | 48,919.35 |
| 6 | 18 | 5,185,581.36 | 27,547.80 | -1,126,026.87 | 49,317.46 |
| 9 | 15 | 5,187,190.16 | 29,156.60 | -1,126,377.52 | 49,668.11 |
| 5 | 15 | 5,190,998.16 | 32,964.60 | -1,127,204.41 | 50,495.01 |
| 15 | 13 | 5,191,792.45 | 33,758.89 | -1,127,377.76 | 50,668.36 |
| 13 | 11 | 5,197,159.46 | 39,125.90 | -1,128,544.06 | 51,834.66 |

Table S4: Estimated demographic parameters for the Spanish, Chinese and Scandinavian groups of *Pinus sylvestris* based on the models 3 and 7. 2.5% and 97.5% are estimated 95% confidence interval values, calculated only for the model 3. $N1_S$, $N1_C$, $N1_N$ are the current effective population sizes of the Spanish, Chinese and Scandinavian groups, respectively. $N2_S$, $N2_C$, $N2_N$ are the historic effective population size of the Spanish, Chinese and Scandinavian groups, respectively. $N_{SPLIT}$ is the effective population size of the ancestor of the Chinese and Scandinavian groups for the model 3, and the effective population size of the ancestor of the Spanish and Scandinavian groups for the model 7; $N_{ANC}$ is the effective population size of the ancestor of all three groups. $Mn_{i \rightarrow j}$ is the population migration rate from group i to group j expressed in number of individuals per generation, n=1 represents the current migration events, n=2 represents the historic migration events. For example, $M1_{S \rightarrow C}$ represents the current migration rate from the Spanish group to the Chinese group. $M_{S \rightarrow SPLIT}$ is the migration rate from the Spanish group to the ancestor of the Chinese and Scandinavian groups; $M_{SPLIT \rightarrow S}$ is the migration rate from the ancestor of the Chinese and Scandinavian groups to the Spanish group. $M_{C \rightarrow SPLIT}$ is the migration rate from the Chinese group to the ancestor of the Spanish and Scandinavian groups; $M_{SPLIT \rightarrow C}$ is the migration rate from the ancestor of the Spanish and Scandinavian groups to the Chinese group. $T_1$ is the time of the population size changes within each group; $T_2$ is the split time between the Chinese and Scandinavian groups for the model 3 and the split time between the Spanish and Scandinavian groups for the model 7; $T_3$ is the split time between the Spanish group and the ancestor of the Chinese and Scandinavian groups for the model 3 and the split time between the Chinese group and the ancestor of the Spanish and Scandinavian groups for the model 7.

| | Model 3 | | | Model 7 |
|---|---|---|---|---|
| Parameter | Value | 2.50% | 97.50% | Value |
| $N1_S$ | $3.76\times10^5$ | $2.84\times10^5$ | $7.79\times10^5$ | $9.97\times10^5$ |
| $N1_C$ | $6.00\times10^5$ | $4.01\times10^5$ | $1.10\times10^6$ | $6.60\times10^5$ |
| $N1_N$ | $9.20\times10^5$ | $7.26\times10^5$ | $1.14\times10^6$ | $2.40\times10^5$ |
| $N2_S$ | $1.88\times10^5$ | $1.44\times10^5$ | $1.90\times10^5$ | $8.18\times10^5$ |
| $N2_C$ | $7.77\times10^4$ | $5.06\times10^4$ | $8.42\times10^4$ | $7.77\times10^4$ |
| $N2_N$ | $3.36\times10^5$ | $2.55\times10^5$ | $3.75\times10^5$ | $1.25\times10^5$ |
| $N_{SPLIT}$ | $4.36\times10^4$ | $1.08\times10^4$ | $5.68\times10^4$ | $8.24\times10^4$ |
| $N_{ANC}$ | $9.50\times10^3$ | $9.15\times10^3$ | $1.48\times10^4$ | $4.89\times10^3$ |
| $M1_{C\rightarrow S}$ | 29.50 | 14.25 | 90.71 | 125.08 |
| $M1_{N\rightarrow S}$ | 114.92 | 73.40 | 380.83 | 121.92 |
| $M1_{S\rightarrow C}$ | 71.20 | 34.79 | 240.72 | 134.91 |
| $M1_{N\rightarrow C}$ | 211.08 | 100.09 | 651.17 | 69.72 |
| $M1_{S\rightarrow N}$ | 134.46 | 82.98 | 305.89 | 71.17 |
| $M1_{C\rightarrow N}$ | 170.65 | 97.33 | 366.68 | 40.50 |
| $M2_{C\rightarrow S}$ | $7.15\times10^{-1}$ | $5.69\times10^{-4}$ | 4.38 | 0.01 |
| $M2_{N\rightarrow S}$ | $3.29\times10^{-4}$ | $2.84\times10^{-4}$ | 5.52 | 0.32 |
| $M2_{S\rightarrow C}$ | $2.45\times10^{-1}$ | $2.19\times10^{-4}$ | 1.09 | 0.00 |
| $M2_{N\rightarrow C}$ | $5.35\times10^{-1}$ | $3.66\times10^{-4}$ | 5.29 | 5.91 |
| $M2_{S\rightarrow N}$ | $1.06\times10^{-1}$ | $2.96\times10^{-4}$ | 2.82 | 0.00 |
| $M2_{C\rightarrow N}$ | $3.39\times10^{-1}$ | $7.55\times10^{-4}$ | 6.31 | 11.21 |
| $M_{SPLIT\rightarrow S}$ | $3.19\times10^{-2}$ | $1.09\times10^{-3}$ | 10.34 | |
| $M_{S\rightarrow SPLIT}$ | 13.36 | 10.18 | 18.67 | |
| $M_{C\rightarrow SPLIT}$ | | | | 5.01 |
| $M_{SPLIT\rightarrow C}$ | | | | 0.02 |
| $T_1$(KYA) | 60.75 | 27.95 | 88.07 | 58.4 |
| $T_2$(MYA) | 0.57 | 0.49 | 0.63 | 0.93 |
| $T_3$(MYA) | 6.20 | 5.17 | 6.20 | 6.98 |

Table S5: The retained environmental variables with low correlation from the gradient forest analysis, ordered by importance. These variables were then used as input for the redundancy analysis and BayPass. Variables in bold were retained in the RDA analysis after the forward selection step.

| **Name** | **Importance ($R^2$)** | **Variable** |
|---|---|---|
| **bio4** | 0.00807 | Standard deviation of temperature seasonality |
| clay | 0.00213 | Proportion of clay particles in the fine earth fraction (0–2 µm) (g/kg) |
| uvb4 | 0.00212 | Mean UV-B of lowest month |
| cfvo | 0.00206 | Volumetric fraction of coarse fragments (cm3/dm3) |
| **gst** | 0.00194 | Mean temperature of all growing season days based on TREELIM |
| silt | 0.00172 | Proportion of silt particles in the fine earth fraction (2–50 µm) (g/kg) |
| **bio8** | 0.00172 | Mean daily temperature of the wettest quarter |
| **cec** | 0.00168 | Cation exchange capacity of the soil [mmol(c)/kg] |
| **bio15** | 0.00167 | Precipitation Seasonality (Coefficient of Variation) |
| **elev** | 0.00158 | Elevation values |
| **bio18** | 0.00154 | Precipitation of the warmest quarter |
| phh2o | 0.00140 | Soil pH (pH x 10) |
| **gdd** | 0.00135 | Growing degree days |
| **nitrogen** | 0.00114 | Total nitrogen (N) (g/kg) |
| ocd | 0.00114 | Organic carbon density ($kg/m^3$) |
| consensus1 | 0.00109 | Evergreen/deciduous needleleaf trees |
| bio12 | 0.00102 | Annual precipitation |
| **ai** | 0.00101 | Arid index |
| gdd5 | 0.00087 | Heat sum of growing degree days above 5°C temperature accumulated over 1 year |
| consensus4 | 0.00082 | Mixed/other trees |
| kg5 | 0.00046 | Köppen-Geiger climate classification according to Troll and Paffen (1964) |

Table S6: Environmental variables (COV) significantly associated with population allele frequencies sorted by outlier significance. Rows in bold indicate outlier loci which are deemed as significantly differentiated ($X^tX$) at $-\log_{10}$(*p*-value) > 3 (*p*-value <0.001). SNP coordinates are given for the *P. taeda* v.1.01 genome and annotations were extracted from that genome (Neale *et al.*, 2014; Zimin *et al.*, 2014). *β* is the posterior mean effect size estimate, expressed in the unit of standard deviations after scaling of all the COV to a unit variance. *β*, its standard deviation *σ(β)*, the posterior inclusion factor (PIP) and the Bayes factor in deciban (BF dB) are from the multivariate analysis[B] if significant, otherwise from the individual analysis[A].

| COV | SNP coordinate | *β* | *σ(β)* | PIP | BF dB | -log10(*p*-value($X^tX$)) | annotation |
|---|---|---|---|---|---|---|---|
| **bio8[A], ai[AB]** | **tscaffold2950:173186*** | **-0.073** | **0.010** | **1** | **62.97** | **7.79** | **Phenazine biosynthesis PhzF protein** |
| **bio8[A], ai[AB]** | **tscaffold2950:173338*** | **-0.068** | **0.010** | **1** | **62.97** | **7.68** | **Phenazine biosynthesis PhzF protein** |
| **bio8[A], ai[AB]** | **scaffold10772.1:113205** | **-0.052** | **0.028** | **0.81** | **26.15** | **5.22** | |
| **ai[AB]** | **tscaffold9099:303439*** | **-0.028** | **0.025** | **0.58** | **21.42** | **5.11** | |
| **bio8[A], ai[AB]** | **tscaffold7790:294517*** | **-0.061** | **0.011** | **0.9999** | **59.96** | **4.68** | |
| **bio8[A], ai[AB]** | **tscaffold7790:294447*** | **-0.061** | **0.011** | **1** | **62.97** | **4.66** | |
| **bio8[A], ai[AB]** | **tscaffold6844:269565*** | **-0.060** | **0.009** | **1** | **62.97** | **4.46** | **Signal peptide peptidase-like 2** |
| **bio8[A], ai[AB]** | **tscaffold7790:281806** | **-0.060** | **0.011** | **0.9998** | **56.95** | **4.31** | |
| **ai[AB]** | **tscaffold694:220164** | **-0.059** | **0.011** | **0.9999** | **59.96** | **4.05** | **Cactin-like** |
| **silt[B]** | **scaffold524899.1:139946** | **-0.051** | **0.013** | **0.9983** | **47.64** | **3.47** | |
| **ai[B]** | **tscaffold2950:173172*** | **-0.028** | **0.021** | **0.67** | **22.95** | **3.01** | **Phenazine biosynthesis PhzF protein** |
| bio8[A], ai[AB] | tscaffold2950:173334* | -0.044 | 0.009 | 0.9985 | 48.19 | 2.98 | Phenazine biosynthesis PhzF protein |
| ai[AB] | tscaffold2950:173208 | -0.036 | 0.009 | 0.99 | 40.89 | 2.92 | Phenazine biosynthesis PhzF protein |
| ai[AB] | scaffold609862.1:89176 | -0.055 | 0.010 | 0.99 | 42.61 | 2.24 | Signal peptide peptidase-like 2 |
| ai[AB] | scaffold653776.3:196917 | -0.041 | 0.026 | 0.73 | 24.37 | 2.11 | |
| consensus4[A] | tscaffold170:80069 | -0.030 | 0.021 | 0.73 | 24.17 | 2.00 | |
| consensus4[A] | tscaffold170:79899 | -0.029 | 0.020 | 0.72 | 24.12 | 1.88 | |
| ai[AB] | scaffold56952.1:75933 | -0.033 | 0.029 | 0.59 | 21.59 | 1.76 | |
| ai[A] | scaffold56952.1:75934 | -0.041 | 0.027 | 0.72 | 24.12 | 1.74 | |
| consensus4[A] | tscaffold170:80054 | -0.022 | 0.020 | 0.59 | 21.56 | 1.69 | |
| consensus4[B] | scaffold757680:9102 | 0.046 | 0.004 | 1 | 62.97 | 1.61 | Non annotated CDS |
| silt[B] | tscaffold5963:28147 | 0.029 | 0.005 | 0.998 | 46.94 | 1.58 | |
| consensus4[A] | tscaffold170:80039 | 0.023 | 0.022 | 0.57 | 21.10 | 1.55 | |
| silt[B] | C32426610:11265 | -0.035 | 0.019 | 0.81 | 26.29 | 0.78 | |
| silt[B] | scaffold895545:68633 | 0.048 | 0.004 | 1 | 62.97 | 0.52 | |
| silt[B] | tscaffold6158:46120 | -0.025 | 0.007 | 0.95 | 32.34 | 0.52 | Mannosyl-oligosaccharide 1,2-alpha-mannosidase MNS1-like isoform X1 |
| consensus4[B] | C32539084:34709 | -0.020 | 0.012 | 0.75 | 24.75 | 0.47 | |
| silt[B] | scaffold467754.1:43814 | -0.037 | 0.004 | 1 | 62.97 | 0.37 | Probable xyloglucan endotransglucosylase/hydrolase protein 23-like |
| silt[AB] | scaffold39803.3:5126 | -0.025 | 0.019 | 0.66 | 22.77 | 0.27 | |
| cfvo[A] | tscaffold4682:859392 | -0.027 | 0.015 | 0.80 | 26.04 | 0.23 | Ankyrin repeat-containing protein |

| COV | SNP coordinate | *β* | *σ(β)* | PIP | BF dB | -log10(*p*-value($X^tX$)) | annotation |
|---|---|---|---|---|---|---|---|
| cfvo[AB] | tscaffold4682:859528 | -0.026 | 0.012 | 0.87 | 28.07 | 0.21 | Ankyrin repeat-containing protein |
| cfvo[AB] | tscaffold4682:859594 | -0.029 | 0.008 | 0.98 | 36.36 | 0.20 | Ankyrin repeat-containing protein |
| cfvo[AB] | tscaffold4682:859500 | -0.027 | 0.010 | 0.92 | 30.39 | 0.19 | Ankyrin repeat-containing protein |
| cfvo[AB] | tscaffold4682:859527 | -0.023 | 0.012 | 0.83 | 26.81 | 0.19 | Ankyrin repeat-containing protein |
| silt[B] | scaffold859842:2650 | 0.027 | 0.006 | 0.98 | 37.11 | 0.17 | |
| cfvo[A] | tscaffold4682:859387 | -0.022 | 0.016 | 0.69 | 23.46 | 0.18 | Ankyrin repeat-containing protein |
| cfvo[AB] | tscaffold4682:859582 | -0.026 | 0.012 | 0.87 | 28.36 | 0.17 | Ankyrin repeat-containing protein |
| silt[B] | scaffold892452.1:273283 | 0.017 | 0.010 | 0.77 | 25.17 | 0.13 | Ribosomal RNA-processing protein 7 homolog A-like |
| ai[B] | scaffold387992:151165 | -0.012 | 0.011 | 0.57 | 21.12 | 0.10 | |
| silt[A] | scaffold572756.1:27700 | -0.009 | 0.009 | 0.55 | 20.80 | 0.07 | |
| silt[B] | scaffold76324:25187 | -0.012 | 0.011 | 0.58 | 21.38 | 0.04 | Pentatricopeptide repeat-containing protein At2g21090-like |

[A] Significant association in the BayPass individual analysis (28), [B] significant in the multivariate analysis (33) and [AB] which were significant in both individual and multivariate analysis (20).
*Outliers in Hall *et al.* (2021).
